# Pollinator abundance shapes sexual selection in an angiosperm

**DOI:** 10.64898/2025.12.10.693606

**Authors:** Estelle Barbot, Mathilde Dufaÿ, Tim Janicke, Benoit Lapeyre, Denis Orcel, François Rousset, Jeanne Tonnabel

## Abstract

Sexual selection is a cornerstone of evolutionary biology potentially operating in all sexually-reproducing organisms. Modern developments in the field revealed that this selective force extends beyond Darwin’s initial focus on access to mates in terms of competition for access to gametes of the other sex. Despite its presumed universality, sexual selection theory remains largely untested in plants compared to animals. This gap may partly stem from challenges in quantifying sexual selection using approaches that account for a critical plant-specific factor: the reliance on pollinators as third-party agents for accessing mates. Here, we quantified sexual selection along consecutive episodes of selection in the hermaphroditic plant *Brassica rapa* by integrating the monitoring of pollinator movements with genetic paternity analyses in experimental populations. Our approach identifies pollen competition for ovules as the primary arena for sexual selection in *B. rapa*. Darwinian competition for access to mates constitutes a secondary force, and was stronger in the male compared to the female sex function, as predicted by classic theory. Importantly, experimentally induced variation in pollinator abundance modulated the balance between pre- and post-pollination sexual selection. Under reduced pollinator abundance, the opportunity for selection on mate acquisition increased. Crucially, we demonstrate that ignoring pollinator movements among plants leads to erroneous quantification of pre-pollination sexual selection. We argue that a unifying theory of sexual selection requires a more comprehensive quantification of pre- and post-pollination episodes of selection, taking into account the specificities of gamete transfer in plants.

## Introduction

Sexual selection is a cornerstone of evolutionary biology, providing a framework to explain the diversity of reproductive strategies observed across the animal kingdom [1,2]. While manifold studies have supported Darwin’s foundational ideas centered on the role of competition for access to sexual partners [3–5], sexual selection has been demonstrated to extend beyond mate acquisition and to encompass post-mating competition for fertilizing gametes [6–9]. However, the vast literature demonstrating the pervasive role of sexual selection in animals contrasts with the limited knowledge on its effects in other sexually reproducing organisms, including flowering plants [10,11]. Conceptual thinking about sexual selection has developed around the core idea that this selective force applies universally to all sexually reproducing organisms [12,13]. A central hypothesis posits that anisogamy leads to a higher fitness return from mate acquisition for the sex that produces the smaller, more abundant gametes (by definition, the male), compared to the sex that produces the larger gametes (the female), and that this difference ultimately makes males more prone to intense competition for access to females and their gametes [12,14]. These sex-differences in the benefit of mate acquisition have been found to prevail across the animal tree of life [3], even though sexual selection in females is potentially more widespread than initially envisioned by the pioneers in the field [15]. Attempts to quantify sexual selection in flowering plants remain scarce, but existing studies have remarkably converged with findings in animals, showing that males obtain stronger fitness benefits from increased access to sexual partners [16–19]. Yet, these first quantitative approaches estimating the strength of sexual selection in angiosperms failed to incorporate the behavior of a plant specific third-party vector – the pollinator, which mediates pollen dispersal in 87.5% of angiosperm species [20]. Integrating pollinator movements between plants into studies on sexual selection in plants is therefore critical to fill this knowledge gap and to disentangle the distinct episodes of selection that occur throughout plant reproduction – from gamete transfer during the pre-pollination phase to ovule fertilization in the post-pollination phase.

In insect-pollinated plants, transport of gametes between sessile sexual partners depends on pollinators such that ecological factors modulating pollinator behavior and abundance are prone to influence the operation of sexual selection. While pollinator abundance has been documented to play a prominent role in plant reproductive biology in terms of seed production or selfing rates [21–25], we have virtually no knowledge on its effect on sex-specific patterns of sexual selection for access to mates. This gap contrasts with the abundant animal literature that demonstrates the role of ecological factors, such as population density, on the strength and direction of sexual selection [26–31]. We hypothesized that variation in pollinator abundance is likely to alter the relative importance of pre- and post-pollination episodes of sexual selection by affecting both the baseline probability of accessing a mate and the level of competition among pollen donors once mates have been accessed. At low pollinator abundance, both sexual functions become more limited by the reduced transport of pollen between mates, which is expected to heighten the importance of the pre-pollination episode of sexual selection to access more mates [32]. Conversely, a high pollinator abundance may intensify post-pollination mechanisms by increasing the quantity and diversity of pollen deposited on stigmas, leading to greater competition among deposited pollen from different donors for fertilizing ovules [33]. The combination of higher pollen loads and stronger polyandry could lead to intensified filtering of more competitive pollen donors through male-male competition and/or cryptic female choice. Previous attempts at quantifying sexual selection in plants have relied exclusively on paternity analyses to estimate the number of sexual partners [16–19], which precludes the ability to disentangle the contribution of accessing mates vs. ovules and therefore confounds the outcome of pre- and post-pollination episodes of sexual selection [34]. To test our hypothesis on the effect of pollinator abundance on the relative importance of pre- and post-pollination sexual selection, it is essential to estimate the number of mates accessed through pollinator movements independently from the ability of pollen donors to fertilize ovules.

We quantified sexual selection at key steps of reproduction in an insect-pollinated plant and tested experimentally how pollinator abundance affects the operation of sexual selection at both pre- and post-pollination episodes in each sex function of the hermaphroditic plant *Brassica rapa* pollinated by its main pollinator *Bombus terrestris [22,35,36]*. To decompose sexual selection episodes, we performed exhaustive monitoring of pollinator movements at the flower scale in 16 experimental populations exposed to three levels of pollinator abundance and genotyped the resulting seeds for paternity assignment. The monitoring of pollinator movements enabled the estimation of a key variable in the quantification of sexual selection – the observational number of mates – defined as the number of sexual partners of a focal plant, accessed in the pre-pollination phase by pollinators coming from the focal plant in the male role, or going to the focal plant in the female role. As the inferred number of mates depends on assumptions about pollen carry-over (i.e. the number of successive flower visits during which pollen export remains effective), we explored the robustness of our results with respect to such assumptions. Paternity analyses of the produced seeds provided critical insights into the post-pollination episode of selection by measuring fertilization success, which is the outcome of the filtering of pollen donors once pollen is deposited on pistils. Interactions between mates are arguably more difficult to estimate in plants compared to animals because a single pollinator visit does not guarantee effective pollen transfer to the pollinator, its transport to a potential mate, and its deposition on a mate’s stigma. Because a simple count of the observational number of mates does not capture these complexities, we quantified two complementary measures: (i) the average number of flowers visited at least once on mates by pollinators visits between the focal plant and its mates (flowers “visited on mates” hereafter), and (ii) the average number of flowers visited on the focal plant itself. The first measure reflects, for the male function, the mean number of flowers (i.e., competitive arenas) on which a plant’s pollen competes, and for the female function, the average quantity of pollen received from mates. The second measure captures, for the male function, the extent of pollen export from the focal plant, and for the female function, the number of flowers that received pollen. The number of flowers visited on mates also plays a central role in our analyses to account for another plant’s characteristic: the simultaneous production of several reproductive organs associated with their modular nature. In particular, it allowed us to quantify the relative importance of reaching more mates vs. reaching more of their flowers in episodes of sexual selection. We finally tested the effects of pollinator abundance on standardized metrics of sexual selection [12,37–41]. Integrating observational and genetic data in our study enabled us to disentangle the relative importance of independent and successive episodes of sexual selection in a plant.

## Results

### Characterization of sexual selection in an insect-pollinated plant

Decomposing male reproductive success into components depicting distinct episodes of sexual selection has become a classical approach to quantifying their relative importance [38,42]. We extended this approach by integrating observational data on pollinator visits with genetic data to evaluate the relative contributions of selection on access to mates (pre-pollination) vs. fertilization of ovules (post-pollination). Specifically, we partitioned variance in male reproductive success into components reflecting the pollen donor’s ability to acquire observational mates, to reach more flowers per mate, to fertilize a greater proportion of seeds, and to access flowers with higher seed production (Fig. 1A). Our results revealed that the primary driver of variance in male reproductive success was seed fertilization ability - a fitness component typically associated with post-pollination performance (Fig. 1A, Table S1). In contrast, the observational number of sexual partners, and especially the number of flowers visited on mates, contributed only little to variance in male reproductive success (Fig. 1A). Beyond the fact that not all visited flowers develop into fruits (63.8% ± 19.7 SD; Fig. S1), the potential fitness gain of visiting more flowers per mate could be offset by less and less pollen deposited on stigma along the flower visit sequence. Pollen donors that concentrate on few flowers per mate could then be over-represented and thus win pollen competition, unlike those that distribute their pollen over a larger number of flowers.

**Fig. 1.**
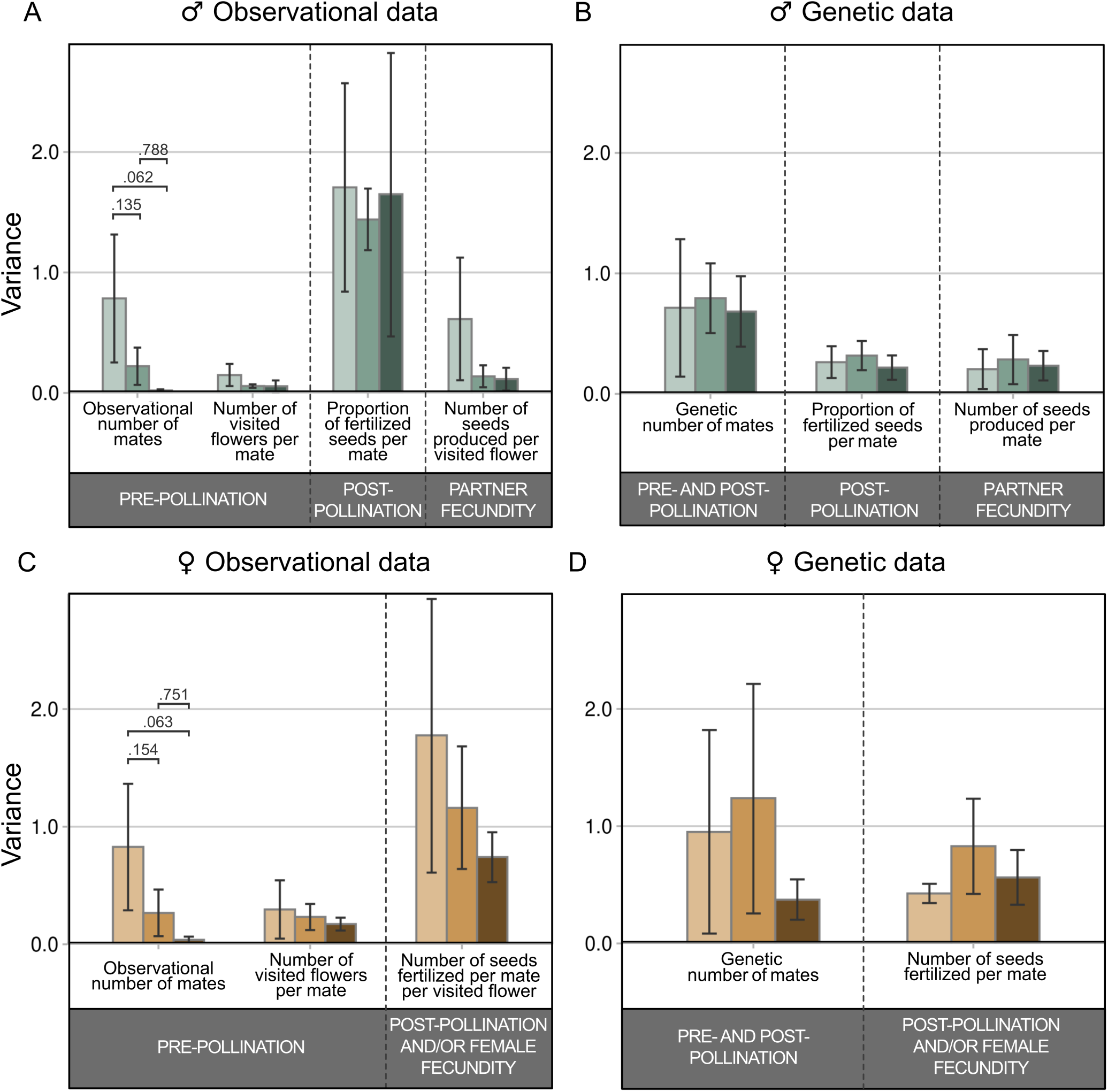
Components of sexual selection across pre- and post-pollination episodes for male and female sexual function in *B. rapa* using observational and genetic data. Variance in outcrossed reproductive success (mean ± 95% bootstrap confidence intervals) for the male (**A, B**) and female (**C, D**) sexual function is decomposed using either the combination of observational and genetic data (**A, C**) or solely genetic data (**B, D**) for the same experimental populations of *B. rapa* exposed to varying pollinator abundance (low: light color; medium: medium color; high: dark color). Significant overall differences in variance components between pollinator treatments, as revealed by likelihood ratio tests on nested models (Table S3), are indicated in the figure by black connecting lines with corresponding adjusted p-values obtained from pairwise comparisons of treatment modalities using post-hoc Tukey’s HSD tests.

The alternative, traditionally used approach, which relies solely on data from genetic parentage analysis to estimate pre-pollination mating success [17,28,30,43] partitions variance into components reflecting a pollen donor’s ability to acquire genetic mates, to fertilize a greater proportion of seeds, and to access mates with higher fertility. This decomposition led to misleading conclusions about the relative importance of pre-vs. post-pollination episodes of selection compared to our pollinator-based approach as it identified the number of sexual partners, instead of seed fertilization, as the primary driver of variance in male reproductive success (Fig. 1B, Table S1). For female reproductive success, the decomposition of variance based on observational data identified a smaller contribution of pre-pollination components of fitness, including both the number of mates and the number of visited flowers per mate, compared to the number of seeds fertilized by mates, a variable capturing mainly female fecundity (Fig. 1C, Table S1). In contrast, variance decomposition based on genetic data alone suggested roughly equivalent contributions of all variance components (Fig. 1D). Our pollinator-explicit approach for the decomposition of episodes of sexual selection highlights the post-pollination phase as the predominant arena driving variance in plant fitness through the male sexual function in *B. rapa*, in stark contrast to results based solely on genetic parentage, which inherently conflate pre- and post-pollination selection episodes [44].

Despite the prominent importance of the post-pollination episode of selection in *B. rapa*, we found support for Darwinian sex roles in terms of stronger pre-pollination sexual selection in the male sexual function as inferred from Bateman gradients (i.e. the slope of the linear regression of offspring number on the observational number of mates, computed for individuals that were visited by a pollinator at least once; 12, 35). Male reproductive success was positively correlated with the observational number of mates at both medium and high pollinator abundances (Fig. 2A; Table S2). This translated into a sex-difference in fitness benefits associated with the observational number of mates (Fig. 2A, Table S2). Given that the contribution of variance in observational number of mates on fitness declined with pollinator abundance (Fig. 1A, Table S3), the scope for selection on the male sex function on mate acquisition in *B. rapa* might be restricted to conditions of intermediate pollinator abundance. In contrast to this sex-specific pattern, both male and female reproduction increased with the number of flowers visited both on mates (i.e. a Bateman gradient computed on another component of pre-pollination performance; Fig. 2B) and on the plant itself (Fig. 2C) at the higher pollinator abundance. Female reproductive success increased with larger pollen loads, but this was primarily due to the production of more selfed seeds (Fig. S2, Table S2). Indeed, the effect of pollen quantity on female reproduction was not consistent when comparing total seed production to strictly outcrossed seed production (Fig. S2). Again, analyses based on genetic rather than observational number of mates erroneously suggested that increased number of genetic mates was beneficial in both sexual functions (Fig. 2D). Male and female reproductive success were not correlated whereas the observational number of mating partners acquired through the male and female function were positively correlated in the medium and high pollinator abundance treatments (Table S4).

**Fig. 2.**
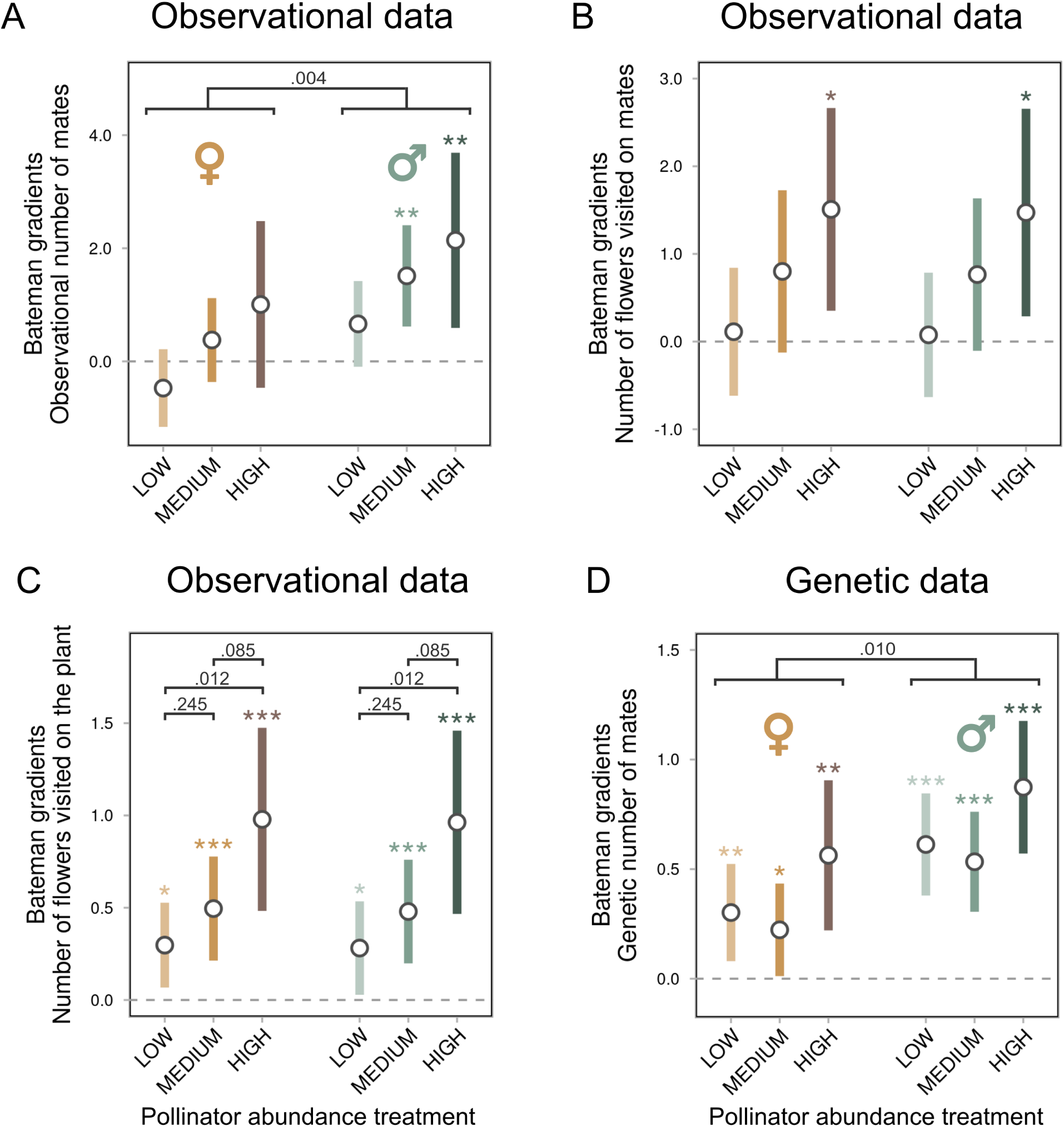
Fitness benefits of mate acquisition and pollinator visitation in the pre-pollination phase for both sexual functions across varying pollinator abundances (low: light color; medium: medium color; high: dark color). Bateman gradients are computed using total reproductive success (estimate ± 95% confidence intervals) based on the observational number of mates (**A**), the number of flowers visited on mates (**B**), the number of flowers visited on the plant itself (**C**), and the genetic number of mates (**D**). Variables related to the female function are represented in shades of orange, while those related to the male function are depicted in shades of green. Bateman gradients significantly different from zero are indicated above each bar in the corresponding color, with asterisks denoting p-value obtained from t-tests (• < .10, * < .05, ** < .01, *** < .001). Significant effects of treatment or sex, identified via likelihood ratio tests (Table S2), are indicated by black lines marking pairwise comparisons performed with Tukey tests and annotated with the respective significance.

### Effect of pollinator abundance on sexual selection

As expected, our manipulation of pollinator abundance led to corresponding changes in the number of pollinator visits received by plants, with a significant positive effect of pollinator abundance on this variable (Fig. S3A; Table S5). Our exhaustive monitoring of pollinator behavior revealed that those changes in pollinator visits translated into increased observational number of mates reached through pollinator movements by the male function and received by the female function (Fig. 3A). Increased pollinator abundance however did not affect the quantity of pollen deposited on stigmas (Fig. S3B) nor plant reproductive success (Fig. S3C) nor the number of flowers visited both on mates (Fig. S3D) and on the plant itself (Fig. S3E).

**Fig. 3.**
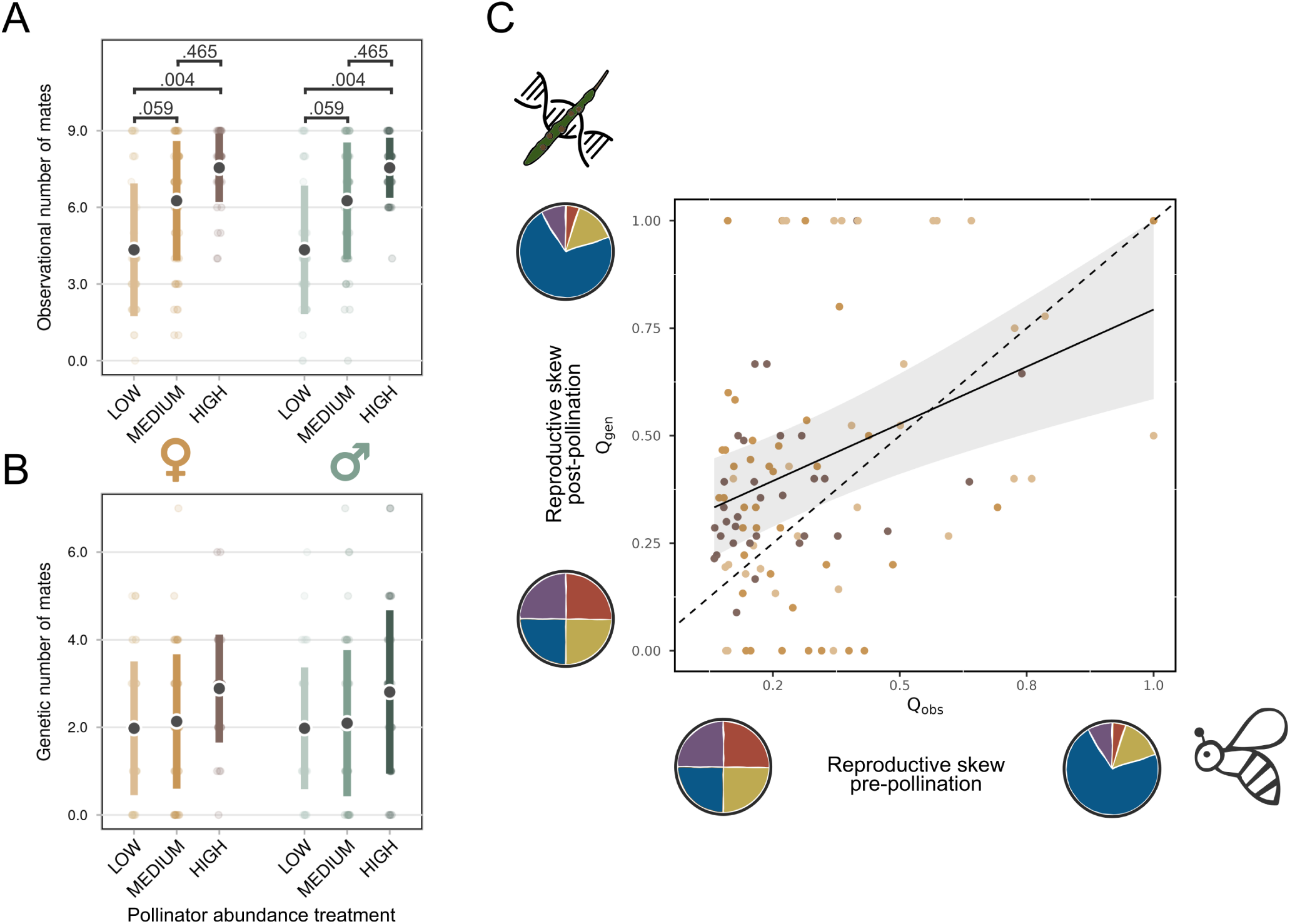
Filtering of mates during the post-pollination phase according to pollen donor diversity during the pre-pollination phase. The effect of pollinator abundance (low: light color; medium: medium color; high: dark color) on the observed number of mates during the pre-pollination episode (**A**) contrasts with the lack of a treatment effect on the number of mates that successfully fertilized seeds after the post-pollination episode (**B**). Regression of pre-pollination reproductive skew (Q_obs_) against post-pollination reproductive skew (Q_gen_) is shown by the solid line with the associated 95% confidence interval in grey (**C**). The relationship significantly deviated from a 1:1 line (dashed), indicating that both low and high pre-pollination skews among pollen donors were homogenized toward intermediate skew levels during the post-pollination phase. Variables related to the female function are represented in shades of orange, while those related to the male function are depicted in shades of green. Significant effects of treatment or sex, identified via likelihood ratio tests (Table S2), are indicated by black lines marking pairwise comparisons performed with Tukey tests and annotated with the respective significance.

In accordance with our hypothesis regarding the relative importance of pre- and post-pollination episodes of selection, pollinator abundance had a negative effect on the opportunity for pre-pollination sexual selection – a classical measure representing the upper limit for selection in a given episode [38]. Specifically, we found that the variance in both the observational number of sexual partners (Fig. 1A, 1C) and in the number of flowers visited on mates (Fig. S4A) significantly decreased for both sexes with pollinator abundance (Table S6). In contrast, the opportunity of selection computed on reproductive success was not affected by pollinator abundance (Fig. S4B). Those results are congruent with the expectation that a decline in pollinator abundance intensifies the opportunity for pre-pollination sexual selection.

Our hypothesis that pollinator abundance strengthens post-pollination sexual selection in the male sex function was not supported by our results as we did not detect any treatment effect on the variance of the proportion of fertilized seeds per mate (Fig. 1A, Table S3). While the observational number of mates increased significantly with pollinator abundance, this increase was no longer significant on the number of genetic mates inferred after both pre- and post-pollination episodes of selection (Fig. 3A and 3B, Table S5). Assuming that our inferences of observational mate numbers accurately reflect pollen deposition, this result might suggest that pollen donors are more effectively filtered out during the post-pollination phase at higher pollinator abundances. To investigate this, we tested whether the difference between reproductive skew before and after pollination - in other words, the inequity in flower contacts by pollen donors across pistils (pre-pollination) versus the inequity in the number of seeds they effectively fertilized (post-pollination) - varied with pollinator abundance. We found no effect of the treatment on the relationship between pre- and post-pollination reproductive skew, nor on the values of the skews themselves (Fig. 3C, Table S7). However, we observed that low pre-pollination reproductive skews (i.e. more equal flower contact among pollen donors) translated into higher post-pollination skews (i.e. less equal fertilization success), and vice versa (Fig. 3C, Table S7). This pattern suggests that when pollinator visits are uneven among pollen donors, the overrepresented donors tend to be filtered out during post-pollination processes. In contrast, when pollen donors contribute more equally in the pre-pollination phase, some are later favored during fertilization (Fig. 3C).

## Discussion

While the role of sexual selection in plants has been acknowledged for decades, the first quantifications in angiosperm species have only recently been conducted [16–19]. Building on these early efforts, which aligned with the vast empirical work performed in animals, our study advances the field by dissecting pre- and post-pollination episodes of selection and by integrating two key plant-specific factors, i.e. the action of pollinators as third-party players in reproduction and the simultaneous production of many reproductive organs. Our results highlight that the ability for fertilizing ovules in the post-pollination phase is a major component of male reproductive success in *B. rapa*. Yet, selection on mate numbers acquired in the pre-pollination phase is operating in the male sexual function especially at intermediate pollinator abundance, effectively validating classical Darwinian sex-roles identified in the animal kingdom [3]. As hypothesized, pollinator abundance affected the relative importance of pre-vs. post-pollination selection, which was driven by a decline in variance in the number of mates acquired during the pre-pollination phase when pollinators became more abundant. Importantly, our study underscores that relying solely on paternity analyses to estimate the number of sexual partners can lead to an overestimation of the selection component that can be attributed to the pre-pollination phase, and to erroneous conclusions regarding the impact of ecological factors, such as pollinator abundance, on the strength and form of sexual selection.

### Sexual selection predominates after pollination

Our quantification of sexual selection across various phases of reproduction led to drastically different conclusions whether the pre- and post-pollination episodes of selection were estimated independently, or not. When sexual partners were counted using genetic paternity, the pollen donor’s ability to access sexual partners emerged as the primary component driving variance in male reproductive success, converging with previous findings in another angiosperm species [17]. In stark contrast, our novel approach, based on the estimation of the actual number of mates obtained in the pre-pollination phase through pollinator movements, identified the pollen donor’s success at fertilising ovules after accessing the mate, as the largest component of sexual selection for the male function. Sexual selection thus acted most prominently in the post-pollination phase in our experimental populations of *B. rapa*, with the pre-pollination phase gaining in importance when the likelihood of exporting pollen to mates diminished due to decline in pollinator abundance. These different conclusions between genetic and observational data regarding which episode of sexual selection is most prominent draws caution on using the genetic number of mates as a metric to quantify sexual selection, as it confounds pre- and post-pollination episodes. Our results concur with previous recommendations in animals [34,44] to independently estimate performance in accessing mates vs. fertilizing gametes when the ultimate goal is to disentangle processes of sexual selection. As previously advocated for animals [44], advancing our understanding of sexual selection in plants will also require detailed characterization of phenotypic traits that affect access to mates and to gametes.

### Male sexual function benefits more from access to mates

Despite the higher opportunity for sexual selection during the post-pollination episode, our study found that sexual selection for increased access to mates during the pre-pollination phase can act on the male function of *B. rapa*. We considered sexual selection to operate in the pre-pollination phase only at intermediate pollinator abundance, as this was the only condition under which the number of observed mates both positively predicted male reproductive success and contributed substantially to variance in male fitness. Importantly, our results support a key prediction of sexual selection theory [12,14]: reproduction via the male function is more dependent on access to mates than reproduction via the female function. These sex-specific fitness returns obtained from mate acquisition align with results of a meta-analysis covering a wide taxonomic breadth in the animal kingdom [3]. It also extends previous work in three angiosperm clades [16–19] by providing the first sex-specific estimates of Bateman gradients in plants, based on direct access of mate numbers through observation of pollinator behaviour. This methodological step forward proved essential for identifying fitness gains obtained from accessing mates, because using genetic rather than observational data resulted in an overestimation of Bateman gradients. This discrepancy is expected for the male function given that genetic data often artificially re-enforces Bateman gradients, as both the number of mates and offspring are estimated through paternity outputs [44]. Importantly, we also observed similar inflation for the female function, highlighting that paternity-based genetic estimates, which cap mate number at the number of viable seeds genotyped, can lead to false positives for female Bateman gradients.

Our study empirically tests whether plant modularity alters the strength and form of pre-pollination sexual selection. We found that the number of flowers visited on mates was a minor component of sexual selection, and amounted to fitness benefits only under high pollinator abundance, where it explained a minimal proportion of the variance in reproductive success. These findings confirm that sexual selection in plants can act through access to mates rather than flowers. Indeed, the fitness benefits of broadly dispersing pollen across many flowers on each mate may be limited – especially if pollen donors that spread pollen over a greater number of flowers per mate experience progressively smaller amounts of pollen deposited on each stigma along the visit sequence [45]. This, in turn, could weaken the probability of ovule fertilization compared to over-represented pollen donors that concentrate on fewer flowers per mate. The reported lesser importance of access to flowers compared to mates in *B. rapa* effectively suggests that the simultaneous production of sexual organs – a common plant feature that may have prevented earlier tests of pre-pollination sexual selection – does not preclude selection for mate access. However, the number of flowers visited, at least once, on a given plant or on its mates may influence the effective transfer of pollen between mates during the pre-pollination phase, thereby potentially shaping all competing pollen donors’ fertilization success in the post-pollination phase. We found that male reproductive success increased with the number of flowers visited on the plant itself, an effect that was amplified under high pollinator abundance. This pattern may indicate a male fitness benefit arising from enhanced pollen export to mates, which could confer post-pollination advantages. However, confirming this hypothesis requires tracking the movement of pollen from multiple donors. Further research exploring the interaction between pre- and post-pollination selection could employ pollen-marking techniques to quantify the relative contribution of competing pollen donors [46,47].

### Pollinator abundance affects patterns of sexual selection

Partially confirming our initial hypothesis, variation in pollinator abundance altered the relative importance of the pre-vs. post-pollination episodes of sexual selection in *B. rapa*. Specifically, variance in the number of sexual partners accessed in the pre-pollination phase increased markedly under low pollinator abundance, suggesting an intensified scope for pre-pollination sexual selection under conditions of pollinator scarcity. Essentially, reduced pollinator abundance led to fewer pollinator visits, which consequently lowered the observational number of mates per plant. Interestingly, under these low pollinator conditions – when plants exhibited greater variation in their observational number of mates – male reproductive success became increasingly influenced by access to highly fecund mates, although this effect did not reach statistical significance. These findings align with classical animal studies demonstrating that ecological conditions can strongly modulate sexual selection processes [26–30,43].

Our results that the opportunity for mate acquisition during the pre-pollination phase increases under reduced pollinator abundance raises the question of whether ongoing pollinator declines could intensify this component of sexual selection, with potential consequences for plant populations. Sexual selection theory posits that this selective force is condition-dependent, favoring individuals in better overall condition who can acquire the resources necessary to develop sexually selected traits [48]. This condition-dependent nature of sexual selection has been well documented in animals [49–51], with possible positive effects on population dynamics [52,53] and the purging of deleterious mutations [54–56]. Whether sexual selection operates similarly in plants - and whether it could lead to unexpected positive effects in the context of pollinator decline - remains an open question. Addressing this requires quantifying genetic correlations between sexually selected traits under low pollinator abundance and other components of plant fitness.

Our initial hypothesis that pollen donors would be increasingly filtered during the post-pollination phase under higher pollinator abundance was not supported by our results. Although the absolute number of mates received by the female function during the pre-pollination phase increased with pollinator abundance, the quantity of pollen deposited on stigmas did not vary significantly. Surprisingly, this rise in polyandry did not lead to a corresponding increase in the number of genetic mates contributing to seed paternity, which may have suggested a treatment-dependent bias in paternity. To investigate whether, consistently with this discrepancy, a larger proportion of pollen donors were filtered out during the post-pollination phase at higher pollinator abundance, we examined whether inequalities in pollen receipt among donors were amplified after pollination. Our results revealed a rebalancing of pollen donor contributions during the post-pollination phase, but which did not vary depending on pollinator abundance. Specifically, when pollen contributions were highly skewed during the pre-pollination phase, initially overrepresented donors tended to be filtered out during post-pollination processes. In contrast, when pollen was more evenly distributed among donors beforehand, some were subsequently favored after pollination. These findings are consistent with our variance decomposition results, underscoring the role of post-pollination sexual selection in reshaping male reproductive skew between pre- and post-pollination phases. Given the apparent importance of the post-pollination episode of selection in this insect-pollinated hermaphroditic species, future research on sexual selection should address whether these filtering processes are strictly attributable to pollen competition or whether a form of cryptic female choice is also involved [33,57].

By accounting for key characteristics of plant reproduction, we provide the most reliable quantification of pre- and post-pollination sexual selection in an insect-pollinated angiosperm. Our independent performance estimates for these two phases uncovered substantial complexities in sexual selection processes that would have remained hidden using genetic data alone. Notably, our findings emphasize that the relative contributions of pre- and post-pollination episodes are not a fixed attribute of a species but are influenced by ecological conditions, particularly pollinator abundance in insect-pollinated plants. We advocate that testing sexual selection theory across anisogamous sexually reproducing clades using approaches directly connected to theory and tailored to the unique characteristics of each clade could foster the development of a unified and comprehensive framework for sexual selection applicable to all sexual organisms.

## Materials and Methods

### Study system and biological materials

Our experimental populations were set up using *Brassica rapa* as the plant species and *Bombus terrestris* as its pollinator. *B. rapa* is a hermaphroditic, herbaceous, insect-pollinated plant species belonging to the Brassicaceae family. It is typically self-incompatible owing to a genetic self-incompatibility locus common to all members of the *Brassica* genus [58,59]. However, this genetic self-incompatibility mechanism is not complete, and variable outcrossing rates are commonly observed in *B. rapa*, including in the line used in this study where selfing has been shown to evolve during experimental evolution [24]. For this study, we used the commercial line *B. rapa* Wisconsin Fast PlantsTM (WFP; Carolina Biological Supply Company, Burlington, North Carolina, USA), a line artificially selected for a short generation time (∼8 weeks) while maintaining substantial phenotypic and genetic variation [60]. The experimental populations in our study were composed of plants originating from five experimental populations of *B. rapa* WFP that had undergone three to four generations of selection under strong polygamy. Specifically, seeds were harvested from 85 parental plants in these populations, where each plant had three flowers pollinated with pollen mixes from 12 anthers collected per plant across the population. After each generation, seeds from the three pollinated flowers were bulked to establish the next generation. The seeds used in this study were mixed from the populations of the third and fourth generations of these populations and are hereafter referred to as the ‘source population’. The pollinator, *B. terrestris*, is a social insect that forms colonies and feeds on a wide variety of plant species [61]. This species is an efficient pollinator of *B. rapa* [22,35,36], making it well-suited for our experimental design.

### Experimental design

We established 16 experimental populations of *our source population of B. rapa*, each exposed to one of three levels of pollinator abundance: low, medium, or high. Seeds from our source population were sown in greenhouses at an experimental platform in Montpellier, France from December, 2021 to January 2022. To ensure sufficient plant availability, seeds were sown in three temporal blocks staggered by one to two weeks, yielding a total of 210 plants (approximately 70 per block). Plants were grown under standardized conditions, with continuous light at 25°C, in separate 0.7 L pots filled with a sterilized soil mixture. Each experimental population consisted of ten flowering *B. rapa* plants randomly chosen among a given temporal block, standardizing plant age to 3-4 weeks at the time of pollination. The selected plants were placed in a mesh cage to confine pollinators during the experiment. Pollinator abundance treatments consisted of exposing the plants to one bumblebee (six replicates), two bumblebees (six replicates), or four bumblebees (four replicates). Prior to each pollination session, pollinators were marked with distinct paint colors to enable individual tracking. Observers recorded verbally the movements of their assigned pollinator in real time, specifying the plant, stem, and flower position visited (by counting flowers downward from the top), for approximately 30 minutes per experimental population, excluding from this counting any initial phase of complete absence of pollinator movements. A flower was only reported as visited if the pollinator came into contact with the sexual organs. To facilitate the recording of pollinator positions, each plant stem had been marked with distinct paint colors, and all open flowers with red paint. Because flower number is known to influence pollinator attractiveness [62], we counted the total number of open flowers on each plant. Flower numbers did not differ between treatments (likelihood ratio tests comparing models with and without pollinator treatment as fixed effect and experimental population as a random effect (low: 15.6 ± 1.02 SE; medium: 14.3 ± 0.95 SE; high: 14.4 ± 1.08 SE; *Χ*^2^=0.18, *P*=0.91, *df* =1, *N* =160). Pollinators were obtained from commercial *B. terrestris* hives (Biobest, Orange, France). To account for potential variability in hive activity, one hive was randomly selected for each pollination session from the four hives purchased with the same representation of each hive across treatments. Across all 16 experimental populations, a total of 160 parental plants were phenotyped and subjected to pollinator observation. After the observation sessions, plants were transferred to an insect-proof greenhouse to ensure controlled fructification. Seeds were subsequently collected for paternity analysis.

### Exhaustive recording of pollinators visits and estimation of observational mate numbers

We transcribed our real-time observational data of pollinator movements recorded at the flower scale for each plant using the BORIS software [63]. This procedure resulted in a dataset documenting, for each visit, the time of the pollinator arrival on a given plant, specifying the plant and flower identity. The primary goal of the pollinator real-time observation was to infer the number of mates obtained by plants during the pre-pollination phase via transport of pollen, referred to in this study as ‘observational number of mates’. This observational mate number was derived solely from the pollinator visit dataset. For the male function, it represents the number of distinct plants to which transfer of a focal plant’s pollen was made possible by pollinator visits. For the female function, it corresponds to the number of distinct pollen donors implied by pollinator visits on the flowers of the focal plants. This estimation of the observational number of mates is conceptually equivalent to the number of copulatory mates traditionally measured in studies of sexual selection in animals [1,39,40,44,64].

While our experiment provides exhaustive data on pollinator movements, we did not track the fate of pollen on pollinator bodies or its transfer among plants. This decision reflects the considerable challenges associated with tracking transport of pollen. Available methods for marking pollen have significant limitations, including: (i) alterations of pollen properties that may hamper its dispersal capabilities [65]; (ii) potential reductions in pollen viability due to chemical treatments, which would bias the decomposition of selective processes occurring in the pre-pollination and post-pollination episodes of selection [47]; and (iii) the limited number of distinguishable colors for marking pollen, which complicates assessments of mate numbers for multiple competing plants [47]. To infer observational mate numbers from pollinator visitation data, we formulated hypotheses regarding the number of successive flower visits during which pollen export remains effective, referred to as pollen carry-over [66,67]. Our primary analysis was based on a pollen carry-over of ten flowers consistent with empirical estimates in a closely related species [68]. To evaluate the robustness of our conclusions, we conducted a sensitivity analysis across a range of pollen carry-over scenarios. Specifically, we tested whether results remained significant for incremental changes in assumed carry-over, and reported the range of values for which our results remained consistent (see Supplementary Tables). The same procedure was applied for the variable quantifying the number of flowers visited on mates as its computation also requires pollen carry-over data (see below).

### Estimation of the intensity of interactions between mates and pollen transfer

Our computation of the observational mate number does not necessarily reflect whether interactions between sexual partners are ultimately effective. For successful pollen transfer between plants – analogous to mating in animals – pollen must be transferred to the pollinator body, transported and deposited onto the stigma of the recipient plant [69]. The actual number of mates accessed may therefore depend on the intensity of pollinator contacts on the mate and the attractiveness of the focal plant. We considered that the number of flowers visited on the mates (averaged across all mates) and on the focal plant may be a reliable indicator of the effectiveness of pollen transfer between mates as it may reflect larger quantities of pollen exchange between mates. In addition to quantifying the intensity of pollinator visits on mates and the focal plant’s attractiveness, we estimated the quantity of pollen deposited on pistils, referred to as pollen loads [70]. Specifically, we collected two to three stigmas per plant five days after the pollination session and once fructification had started, and assessed the number of pollen grains deposited, following the protocol described in the Supplementary Text S1.

### Genotype acquisition, paternity analyses and estimation of the genetic number of mates

When plants had completed their life-cycle, we counted all produced fruits, harvested them, and both counted all seeds for later assessment of the variance in reproductive success via the female function. On average, seed number was assessed in 6.52 (SD ± 4.37) fruits per maternal plant, yielding a total of 959 counted fruits. We assessed germination rate by sampling seeds proportionally to the number of seeds produced (mean ± SD: 22.9 ± 27.4 per maternal plant), which were sown in Petri dishes filled with 40 mL of 10 g/L agar in sterile water (photoperiod: 16:8, temperature: 20-23°C) for one week. For each maternal plant that produced seeds (137 in total), ten seedlings were selected for subsequent DNA extraction and genotyping for paternity analysis, except for 25 individuals that produced fewer than ten viable seeds and for four individuals that failed to produce any viable seeds. DNA extraction, single-step PCR assays, and capillary electrophoresis for both adults and seedlings were performed as described in the Supplementary Text S2. In a nutshell, samples were subjected to a tissue homogenizer, followed by DNA extraction using magnetic beads and an automated purification workstation. Single-step multiplex PCR assays were then used to amplify eight nuclear microsatellites, and electrophoresis was performed with a size standard (Table S8). Alleles at the eight loci were assigned using GeneMapper® (Thermo Fisher Scientific, Waltham, USA) for each parent and offspring, resulting in genotypes for 160 parents and 1,193 seedlings.

Paternity analyses were conducted using CERVUS 3.0.7 [71,72]. This software performs categorical paternity assignments by simulating a large number of reproductive events with simulated parents of known genotypes, sampled accordingly to estimated allelic frequencies in the study population. This simulated dataset with known paternity allows for the calculation of likelihood score distributions and determination of a critical value for assigning the true father to a given seed at a specified confidence interval. Paternity is assigned to a given individual based on likelihood ratios, comparing the likelihood of observing the offspring genotype assuming the individual is the true father, to the likelihood assuming a random male is the father. We used a 80% confidence level (relaxed criterion) to retain the most-likely father, allowing for up to three minimum typed loci and 1% genotyping error (estimated based on mismatches between mothers and offspring, which averaged 0.084% across all eight markers). The mean non-exclusion probability was 0.098 across the 16 sessions, with a range from 0.050 to 0.15. The assignment procedure successfully assigned paternity to 1,099 seedlings (mean ± SD: 6.87 ± 3.97 per maternal plant). Female and male genetic number of mates were computed as the number of sexual partners with whom a given plant shared at least one seed following fertilization. Thus, this variable captures not only access to mating partners determined during pre-pollination selection, but is also affected by fertilization success, which is subject to post-pollination selection. Only a small proportion of the genotyped seeds (7.73%; 85 seeds out of 1099) exhibited genotypes that were not compatible with our observational data of pollinator visits under an infinite carry-over model, confirming the validity of our approach for inferring observational mate numbers based on pollinator observations.

### Estimation of sex-specific reproductive successes

For both sexes, we calculated both total reproductive success and strictly outcrossed reproductive success. Total female reproductive success was quantified as the total number of seeds produced, weighted by the estimated seed germination rate for each female. Outcrossed female reproductive success was estimated as the proportion of seeds produced with another plant (assessed by genotyping seeds) multiplied by total female reproductive success. Male reproductive success was quantified as the sum, across all mothers, of the product of (i) the proportion of seeds sired by the focal male on each mother by (ii) the mother’s reproductive success. Female and male reproductive successes were calculated by either excluding or including the selfed seeds. We also computed reproductive successes without weighting seed production by germination rates to avoid artificial correlation between female reproductive success and the number of genetic partners that may emerge because the genetic number of mates could solely be assessed on germinated seeds for the female function. As most of our results were robust to the inclusion or exclusion of selfed seeds, we presented our results considering total reproductive success but any lack of robustness when using outcrossed reproductive success is shown in supplementary and discussed.

### Statistical analysis

#### Effect of pollinator abundance on pollinator visits and reproductive performance

To test whether pollinator abundance influenced pollinator visits received by individual plants, we first compared mixed-effects models explaining the total number of visits received per plant, with and without treatment as a fixed effect, using likelihood ratio tests. The same procedure was applied to evaluate the effect of pollinator treatment on pollen loads, plant reproductive success, both observational and genetic number of mates, and the number of flowers visited on the mates and on the plant itself. Specifically, we compared nested mixed-effects models explaining sequentially one of these variables and including or not the sex by treatment interaction. Post-hoc Tukey tests were conducted to test for pairwise differences between levels of pollinator abundance and/or sexual functions when significant effects of treatment, sex or their interaction were observed. All models throughout the statistical analysis included random effects for experimental populations and nested random effects for individuals within session, the latter because a plant could be used in different sex roles in different sessions, or because several measures were taken per plant. Each response variable was modeled using the most appropriate error distribution, as reported in Table S5. When appropriate, statistical analyses were performed with and without individuals that failed to reproduce.

#### Opportunity for (sexual) selection based on pollinator abundance

We estimated two classical metrics for quantifying the upper limits of selection and sexual selection: the opportunity for selection (*I*) classically defined as the variance in reproductive success divided by the square of the mean reproductive success, while the opportunity for sexual selection, (*I_S_*) was defined as the variance in mate number divided by the square of the mean mate number (*40*). We estimated *IS* using both observational or genetic mate numbers, as well as on the number of flowers visited on mates and on the plant itself. All computed metrics of selection and sexual selection were calculated for each independent experimental population and each sexual function. Following established recommendations [42,73,74], variances in reproductive success were corrected for the expected binomial sampling error for both sexes. Specifically, variance in male reproductive success was corrected for the binomial sampling of focal offspring, while variance in female reproductive success was corrected for the binomial sampling of outcrossed vs. selfed seeds. The effects of treatment, sex, and their interaction on opportunities for selection and sexual selection were tested using likelihood ratio tests, and differences between pollinator treatment modalities were estimated using t-tests.

#### Quantification of Bateman’s gradients

Bateman gradients classically quantify fitness benefits of increased access to sexual partners [12,39]. Building on classical methods in animals, we estimated Bateman gradients as the slope of the regression of reproductive success on either the number of mates (observational or genetic) or the number of flowers visited on mates and on the plant itself. Individuals that did not reproduce (and thus had zero sexual partner) through either sexual function were removed from these analyses to avoid inflation of the estimated slope. Prior to the estimation of Bateman gradients, we relativized both response and explanatory variables by dividing each value by the mean within each sexual function and experimental population [44,75]. To test whether Bateman gradients differed between sex and pollinator abundance treatments, we built linear mixed models explaining the relativized reproductive success as a function of one of the relativized explanatory variables described above, by pollinator abundance treatment, sex, and their interaction. Specifically, we used likelihood ratio tests to compare models including or not the three-way interaction term. When a significant three-way interaction was detected, we extracted slope estimates and significance levels for each combination of pollinator treatment and sexual function, by modifying contrasts in the model. When the three-way interaction was not significant, we assumed that pollinator treatment effects were consistent between sexual functions and tested for two-way interactions between the explanatory variable and either pollinator treatment or sex. Bateman gradients were calculated using reproductive success estimates that did not account for germination rates. Including germination rates could lead to false detection of positive Bateman gradients in females because the number of genetic sexual partners, by being counted on germinated seedlings, is necessarily positively related to germination rates. Results were also checked by including germination rate. Our quantification of sexual selection was primarily carried out by the traditionally more established metrics in the field, i.e. opportunities of selection and Bateman gradients [12,37]. For the sake of completeness, we also report results obtained for Jones indexes, a metric quantifying the maximal strength of sexual selection as the product of the opportunity for sexual selection (*I_S_)* and the square root of the Bateman gradient [40]. This metric was recently argued to better capture sexual selection in the pre-mating phase in animals [41], and results obtained in our study were congruent with our conclusions based on the other traditional metrics (Table S9).

Both sexual functions are jointly expressed in hermaphrodites, thus resource allocation to acquiring mates in one sexual function could influence the reproductive success of the other [44,73]. Additionally, performance in one sexual function may correlate with performance in the other because individuals with greater resource acquisition can allocate more resources to both sex functions. To describe the reproductive biology of our hermaphroditic species, we first quantified correlations between the two sexual functions in terms of reproductive success, observational and genetic sexual partners. As the production of selfed seeds is necessarily correlated for the male and female sexual functions, this analysis was restricted to the outcrossed fraction of reproductive success. Mixed-effect models were used to regress male-function values on female-function values, accounting for the random effects for experimental populations and nested random effects for individuals within sessions. To complement our primary analysis of Bateman gradients, we estimated cross-sex effects, which capture whether mate numbers obtained via the male function affects reproductive success via the female function, and vice versa [73]. Cross sex-effects were excluded from the main analysis as they were neither statistically significant individually, nor in interaction with sex or treatment (Table S10). Furthermore, their inclusion did not alter the primary results of the analysis.

#### Decomposition of variance in male and female reproductive success

Variance in male and female reproductive success was decomposed into its components, to identify the factors contributing most to the total variance and to investigate how pollinator abundance treatments affected their relative contributions. This variance decomposition was performed using classical methods based on genetic estimates of mate numbers [42], and by adapting this methodology to data on pollinator observation. This allowed us to compare results from genetic mates estimates with those derived strictly from pollinator movements in the pre-pollination episode. Detailed equations for the sex-specific variance decomposition methods, for genetic and observational data, are provided in the Supplementary Text S3. For this specific analysis, we omitted variance in selfed offspring as the main goal was to decompose episodes of sexual selection. In summary, variance in male outcrossed reproductive success was first decomposed into components including the number of genetic partners, the proportion of seeds fertilized per partner, the partner fecundity, and covariances among these factors (Fig. S5). Using observational data, variance in outcrossed male reproductive success was decomposed into the number of observational partners accessed, the number of flowers on which pollinators deposited pollen per mate, the number of seeds produced by mate per flower accessed, the proportion of seeds fertilized within these flowers, and all associated covariances (Fig. S5). For the female function, variance in outcrossed reproductive success was decomposed into the number of genetic mates and the number of offspring produced with each mate, along with the covariance between these two components. Using observational data, variance in outcrossed female reproductive success was also decomposed into the number of mates whose pollen landed on the pollen recipient’s stigma, the number of flowers accessed by pollen of a given mate, and the number of seeds produced by flower accessed. All variance components were computed at the level of the experimental population. Individuals that did not reproduce were included in the computation of the variance in the number of mates. We tested whether these components differed with pollinator abundance, following methods described above for the metrics of opportunities of selection. To test for qualitative differences in the contributions of variance components, we compared nested mixed models with a random effect of the experimental population explaining the variance attributed to each component, with and without the identity of the variance components, using likelihood ratio tests. When variance differences between components were detected, Tukey tests were applied to identify the largest contributing component of reproductive success variance.

#### Inference of filtering by the mother plant

To test for filtering by the mother plant, we compared the diversity of observational mates to the diversity of genetic mates. However, the number of mates, as usually considered in Bateman gradients, is poorly suited for such comparisons. The number of genetic mates of a focal plant in female role was derived from a lower number of observations (the number of genotyped seeds) than the number of observational mates (derived from observations of pollinator visits). So the inferred number of genetic mates is expected to be lower than the number of observational mates, with a reduction factor depending on sample size and on the distribution, among pollen donors, of number of visits on the focal plant, as well as on genetic sample size. To avoid this inherent bias, we used a measure of reproductive skew, defined as the frequency of pairs of distinct visits (or of pairs of parentages of distinct seeds) that involve a single mate, to quantify the diversity of mates: the higher the frequency, the higher the skew. The reduction in genotyped sample size does not *per se* bias the comparison between diversities of observational and retained mates after fertilization. A well-known measure of reproductive skew, Morisita’s index [76,77], is actually identical, up to a multiplicative constant (the number of potential mates), to this frequency (hereafter *Q*).

We thus fitted the frequency *Q_gen_* among genotyped seeds to the frequency *Q_obs_* among observed visits, by linear mixed-effect models. We allowed heterocedasticity of residuals, which is here expected by design, by fitting the residual variance in terms of the number of genotyped seeds from which each response value *Q_gen_* was determined. These heteroscedastic models were adjusted with the `spaMM` R package [78]. The full model allowed for an interaction between *Q_obs_* and pollinator abundance.

In the absence of any filtering by the mother plant, the fit should be consistent with a 1:1 relationship between *Q_gen_* and *Q_obs_*. Diversifying selection by the mother plant will increase *Q_gen_* above *Q_obs_*, and conversely it will reduce *Q_gen_* below *Q_obs_* if the mother plant selects specific mates. We compared more or less constrained models, including a model where the regression line was forced through the point on the 1:1 line corresponding to the highest observed value of *Q_obs_*. If this fit is not rejected by a test in comparison with the unconstrained model, then it cannot be rejected that the *Q_obs_* predicted by the fit are always above or always below *Q_gen_*, depending whether the fitted regression slope of the constrained model is lower or higher than 1, respectively.

## Acknowledgments

Part of the experiments were conducted at the greenhouses of the Centre d’Ecologie Fonctionnelle et Evolutive (CEFE-CNRS), a platform supported by the LabEx CeMEB, an ANR “Investissements d’avenir” program (ANR-10-LABX-04-01). We sincerely thank Marie Challe and Benjamin Viel for their help with pollinator observations in the greenhouse. We are grateful to Maryse Vanderplanck for her advice on beehive management. For their contribution to genetic data acquisition, we thank Élodie Flaven, Fleur Hamoir, and Ulysse Benvegnen. We also appreciate the support of Agnès Mignot, Céline Devaux, and Patrice David regarding helpful discussions on statistical analyses and biological interpretations.

## Funding

This project received funding from:

the European Research Council (ERC) under the European Union’s Horizon 2020 research and innovation program (Grant Agreement No. 101078021; JT)
the University of Montpellier ‘Tremplin ERC’ funding program (project SEXMATE; JT)

## Author contributions

Conceptualization: JT

Resources: DO, BL

Investigation: EB, MD, DO, JT

Formal analyses: EB

Methodology: TJ, FR, JT

Supervision: JT

Writing—original draft: JT, EB

Writing—review & editing: MD, TJ, FR

## Competing interests

Authors declare that they have no competing interests.

## Data and materials availability

All data and code are available in Zenodo (link to be added upon acceptance).

## Supplementary Materials for

### This PDF file includes

Supplementary Text S1 to S3

Figs. S1 to S5

Tables S1 to S10

**Supplementary Text S1.** Pollen loads estimation.

Two to three stigmas were collected on each individual five days after pollination. Our preliminary tests on *B. rapa* Fast Plants showed that harvesting stigmas five days after pollination did not affect seed production. In the current study, the number of seeds produced did not differ between flowers with and without harvested stigmas, as shown by comparing models explaining seed set with or without a variable describing the stigma status (harvest or intact) using likelihood ratio tests (*Χ*^2^=1.43, *df* =1, *N* =590, *P*=.231). This test was performed on a dataset including 590 fruits (i.e. 214 with removed stigmas and 376 with intact stigmas). The observation session and individual plants nested within sessions were treated as random effects in this analysis. Once collected, stigmas were stored in separate eppendorfs containing FAA (ethanol 70%, formaldehyde 35% and acetic acid, 8:1:1), which allows long time conservation of pistil tissues while preserving the integrity of pollen grains before counts. Stigmas were then immersed in a bath of 10µL of sodium hydroxide (NaOH 4 mol.L^-1^) for at least 3 hours to soften tissues, and then washed for one hour in a bath of 10µL of water. Stigmas were then mounted on a microscope slide with Alexander solution to color pollen grains [1], gently crushed to place tissue in the same plane, and observed at x10 magnification (Olympus CX43; Olympus Corporation, Tokyo, Japan). We took pictures of the whole stigmatic surface using a camera adaptator (Olympus U-CMAD3; Olympus Corporation, Tokyo, Japan) for later counts with Image J [2]. To account for potential loss of pollen grains during tissue preparation, we also counted for each sample the remaining pollen grains in the two baths of sodium hydroxide and water by depositing the solutions on Mallassez cells and counting pollen grain directly (2.5µL for each solution). The number of pollen grains, i.e. pollen load, on each stigma was finally obtained by summing counts on all pictures as well as pollen grains counted in the solutions of sodium hydroxide and water.

**Supplementary Text S2.** DNA extraction, PCR and electrophoresis.

Total genomic DNA from adults was extracted from approximately 12 mg of dried leaves, while offspring DNA was directly extracted from young seedlings. To obtain the seedlings, seeds were sown on petri dishes containing agar in sterile water (10 mg·L⁻¹) and incubated for one week under a photoperiod of 16:8 (light:dark) at temperatures ranging from 20 to 23°C.

DNA extraction for both adults and offspring was performed using the NucleoMag Plant kit for DNA purification (Macherey-Nagel Inc., Allentown, USA), following the manufacturer’s instructions. A summary of the extraction process is provided below. Biological samples were initially crushed using a tissue homogenizer with tungsten beads (Qiagen, Hilden, Germany). The samples were then incubated in a thermolyser with RNase and a lysis buffer. After centrifugation, the clear lysates were transferred to deep-well plates (Thermo Fisher Scientific, Waltham, USA) containing a buffer and magnetic beads. DNA purification was automated using a KingFisher® workstation (Thermo Fisher Scientific, Waltham, USA), where successive washes with either kit buffers or 80% ethanol yielded purified DNA, which was eluted in the kit’s elution buffer.

Single-step multiplex PCR assays were performed on each DNA extract to amplify eight nuclear microsatellites (Table S8), using Applied Biosystems™ Master Mix (Thermo Fisher Scientific, Waltham, USA). The PCR cycling program included the following steps: (i) initial denaturation at 95°C for 15 minutes; (ii) amplification for 30 cycles consisting of 95°C for 30 seconds, 58°C (annealing) for 1 minute and 30 seconds, and 72°C for 1 minute; (iii) final extension at 60°C for 30 minutes.

PCR products, diluted 1:120 in distilled water, were analyzed via capillary electrophoresis (Applied Biosystems® 3130 Genetic Analyzer). GeneScan™ 500 LIZ™ (Thermo Fisher Scientific, Waltham, USA) was used as the size standard for allelic profile characterization, which was performed using GeneMapper® software (Thermo Fisher Scientific, Waltham, USA).

**Supplementary Text S3.** Variance decomposition analyses.

Reproductive success was decomposed into its components by extending established methodologies adapted to *Brassica rapa* [3–5] using both the classical metric of the genetic number of mates and the observational number of mates. The latter is employed for the first time in decomposition of plant reproductive success, prompting the development of a novel analytical framework to assess the effects of different components contributing to variance in plant reproductive success. The objectives of this analysis were to: (i) identify the components contributing most to variance in reproductive success for each sexual function, (ii) determine whether the relative contributions of these components varied between pollinator abundance treatments, and (iii) compare results obtained using genetic versus observational mate numbers. To achieve these goals, we first decomposed variance in reproductive success using estimates of mate numbers based on (i) mates sharing at least one produced seed (genetic number of mates, encompassing both pre- and post-pollination processes) or (ii) mates acquired through pollinator movements, which includes only pre-pollination processes. Depending on the proxy used (genetic or observational number of mates), we further decomposed variance into its successive components, as described in detail below.

The analysis focused exclusively on outcrossed reproductive success, as selfing can occur independently of pollinator visits, precluding its decomposition using observational mate numbers. As variance in reproductive success was decomposed at the scale of the population and per sexual function, one population was excluded from the dataset because only two individuals in this population reproduced via outcrossing, precluding us to estimate variance component. Additionally, for variance decomposition using the observational number of mates, we restricted the analysis to outcrossed seeds that were compatible with our data on observed pollinator visits, thereby excluding seeds derived from unobserved pollen transfer, contamination during plant handling, or recording errors. Under an infinite carry-over model, this exclusion resulted in a loss of 11.47% of seeds (85 seeds) and 13.20% of mate pairs. Using a carry-over model with a limit of ten flowers (our biologically realistic hypothesis), seed loss increased to 27.13% (201 seeds), with a 26.73% loss of mate pairs. Note that when we include selfing, the exclusion resulted respectively in a loss of 7.73% of seeds under an infinite carry-over model and of 18.29% of seeds under a carry-over model with a limit of ten flowers. Since bumblebees occasionally exhibit imprecision in their flower visits, we attempted to assign visits to flowers adjacent to those observed, which might account for some unobserved pollen transfer. However, efforts to mitigate data loss by weighting nearby flowers did not effectively resolve these discrepancies. Consequently, we assigned visits strictly to the observed flowers to maintain the dataset’s clarity and avoid unnecessary complexity. The mean error on the sum of the components of the variance of reproductive success compared to the actual variance of reproductive success was 21.32% (± 0.16 SE), all analyses combined.

### Genetic number of mates

- Male function Using the genetic mate number as classically used in variance decomposition, we decomposed male reproductive success as classically performed in previous studies (Equation 1; 3–5)

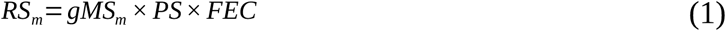

with *gMS_m_* corresponding to the genetic mate number for the male function (encompassing both pre- and post-pollination component) and including individuals with zero mates, *PS* to the average proportion of seeds fertilized per mate (post-pollination component), and *FEC* to the average number of seeds produced by each mate, which represents the partner’s fecundity. Pélissié et al. (2012) provided an approximation of the relativized variance in reproductive success (Equation 2; 5).

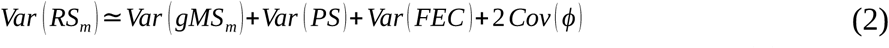

with each variance component being relativized by their grand mean and 2 *Cov* (*ϕ*) representing the sum of the three pairwise covariances between the main components. Results are available in Figure 1B and S5B for the outcrossed reproductive success. Note that only males (and females in the below analyses) with *gMS*> 0 are considered to compute variance in *PS* and *FEC*, variance in these two components are therefore conditional to mate acquisition. Similarly, when focusing on observational mating success, only individuals with *oMS* >0 are considered to compute variance in the subsequent component.
- Female function We decomposed female reproductive success into its different components also following classical methods (Equation 3; 3–5).

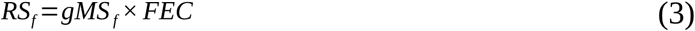

with *gMS_f_* corresponding to the genetic mate number for the female function (encompassing both pre- and post-pollination component) and *FEC* to the average number of seeds fertilized per mate, which represents primarily the female function fecundity and secondarily the partner’s ability to fertilize and develop seeds. We then obtained the formula for the relativized variance decomposition as follows (Equation 4).

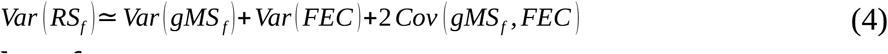

### Observational number of mates

- Male function Using the observational mate number, we developed a novel procedure for decomposing male reproductive success as described in Equation 5.

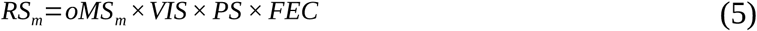

with *oMS_m_* corresponding to the observational mate number for the male function (pre-pollination component) and including individuals with zero mate reached, *VIS* to the average number of visited flowers per mate (pre-pollination component), *PS* to the average proportion of fertilized seeds per mate (post-pollination component, assessed by genotyping seeds), and *FEC* the number of seeds produced per visited flower, which represents the partner’s fecundity. In the variance decomposition performed using observational data, we therefore obtain variance components that correspond strictly to the pre-pollination or to the post-pollination phase, thus enabling the estimation of their relative importance. An approximation of the relativized variance in male reproductive success is then provided in Equation 6:

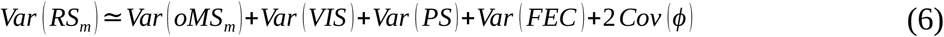

with 2 *Cov* (*ϕ*) representing here the sum of the six pairwise covariances between the main components.
- Female function Using the observational mate number, we developed a novel procedure for decomposing female reproductive success as described in Equation 7.

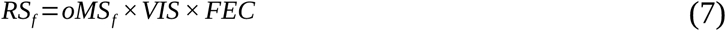

with *oMS_f_* corresponding to the observational mate number for the female function (pre-pollination component), *VIS* to the average number of visited flowers per mate (pre-pollination component) and *FEC* to the average number of seeds fertilized per mate per flower, which represents primarily the female function fecundity and secondarily the partner’s ability to fertilize and develop seeds.

**Fig. S1.**
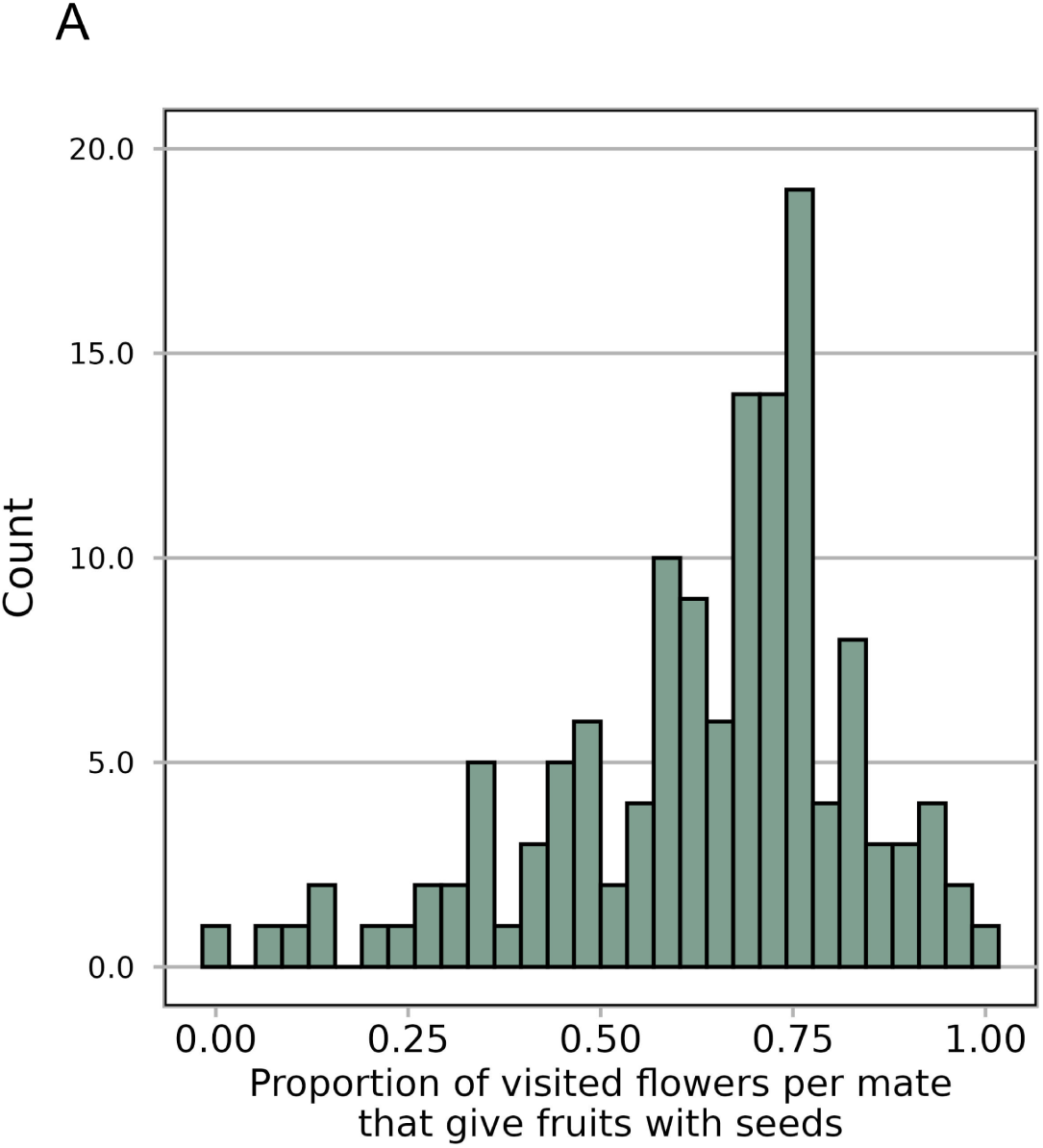
Distribution of the proportion of visited flowers per mate that give fruits with seeds.

**Fig. S2.**
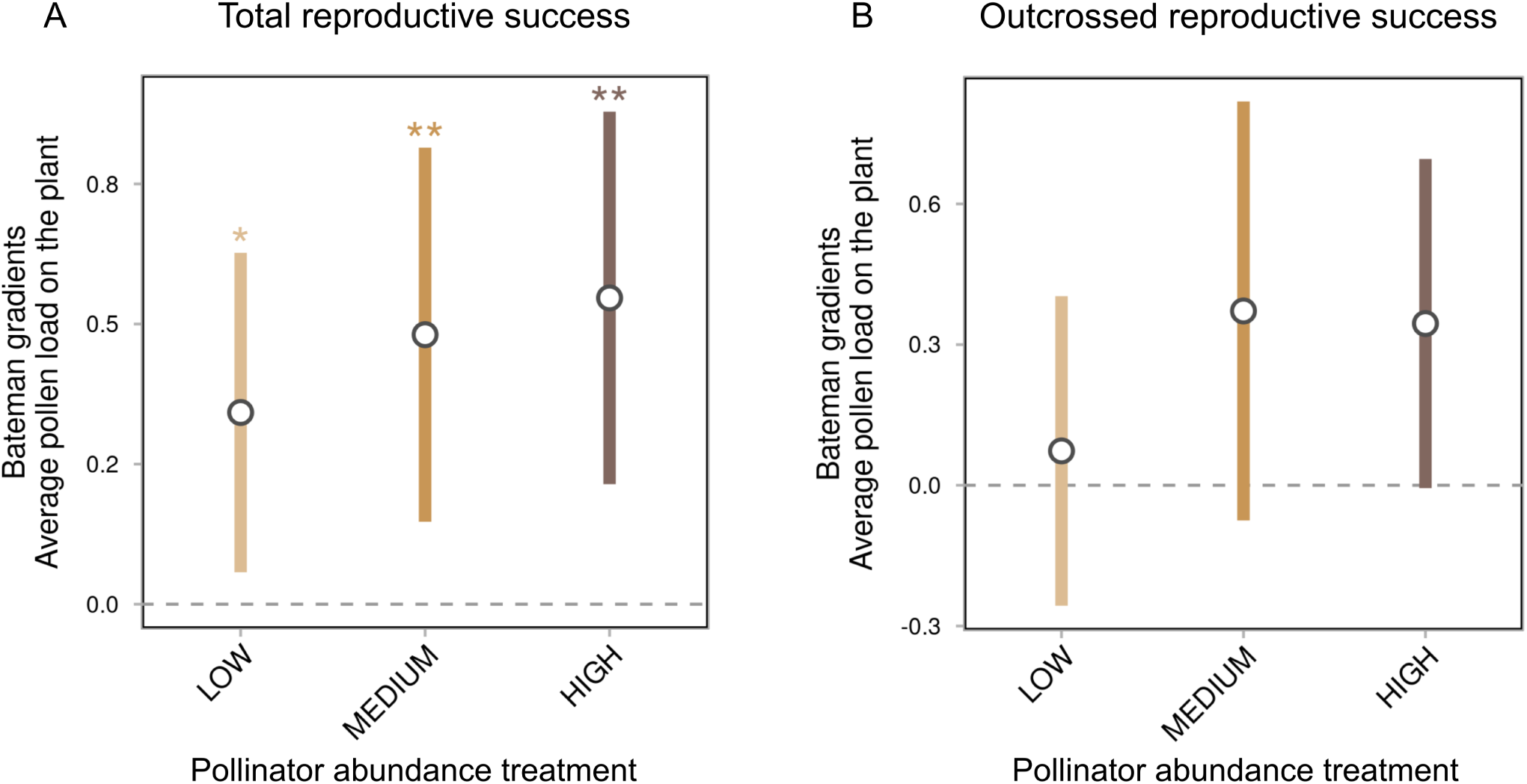
Effects of the pollinator abundance treatment on the Bateman gradients on pollen loads for both reproductive successes. Bateman gradients (estimate ± 95% confidence intervals) on average pollen load on the plant are displayed for each pollinator abundance treatment (low, medium, or high abundance), using total (A) or outcrossed (B) female reproductive success. Bateman gradients differing significantly from zero are indicated above each estimate in the corresponding color, with stars corresponding to the p-value from t-tests (^•^ < .10, * < .05, ** < .01, *** < .001). Significant treatment, sex or interaction effects, as identified by likelihood ratio tests (Table S2), are indicated in this figure by black lines with their associated p-values from t-tests for pairwise comparison between modalities.

**Fig. S3.**
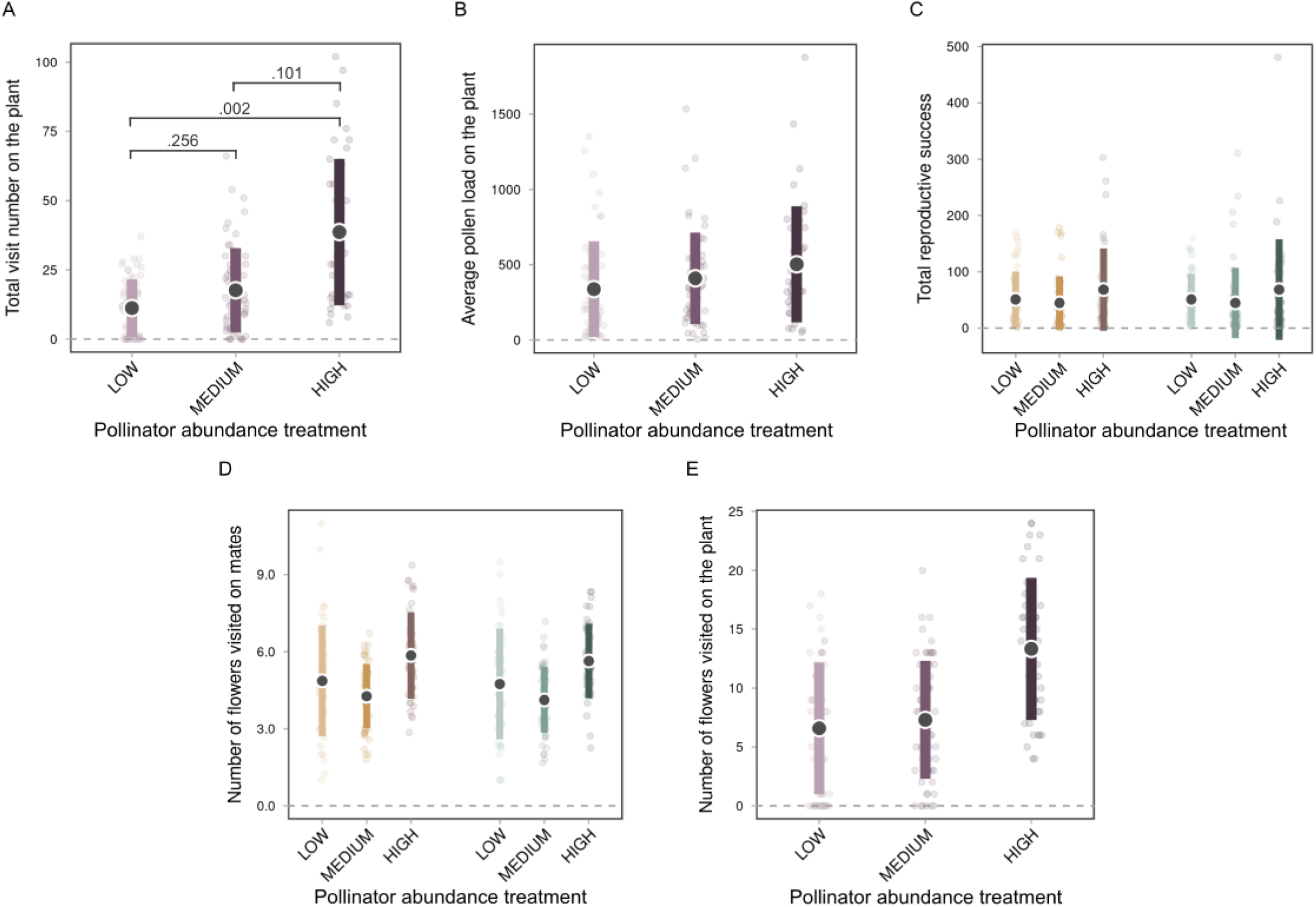
Effect of pollinator abundance on pollinator visitation, pollen loads, and plant reproductive success. The impact of pollinator abundance treatment (low, medium, or high) was assessed on the following variables (mean ± standard deviation): (A) total number of visits to the plant, (B) average pollen load on the plant, (C) total reproductive success, (D) number of flowers visited on mates, and (E) number of flowers visited on the plant. When applicable, variables associated with the female function are shown in shades of orange, those related to the male function in shades of green, and variables shared by both sexual functions in shades of purple. Significant treatment effects, identified by likelihood ratio tests (Table S5), are indicated by p-values from post-hoc Tukey tests for pairwise comparisons between treatment levels shown by black connecting lines.

**Fig. S4.**
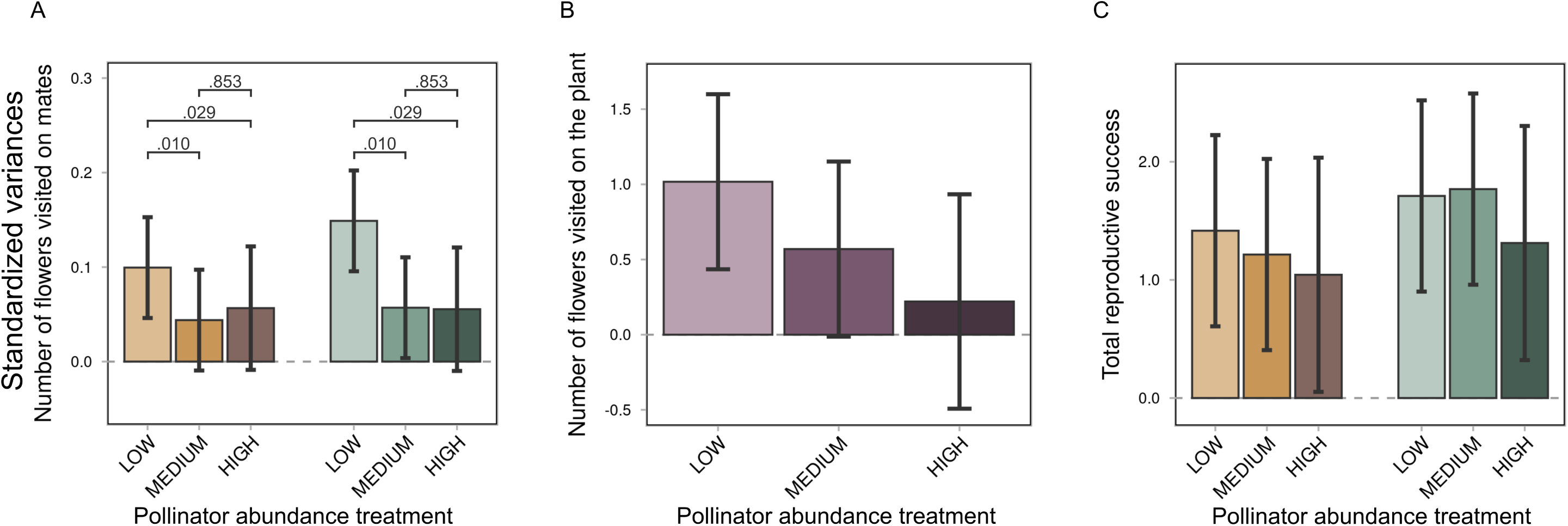
Effects of pollinator abundance on the opportunities for selection. The impact of pollinator abundance treatment (low, medium, or high) was evaluated for the following measures of opportunity for selection (mean ± 95% confidence intervals): (A) number of flowers visited on mates, (B) number of flowers visited on the plant, and (C) total reproductive success. When applicable, variables related to the female function are shown in shades of orange, those related to the male function in shades of green, and variables shared by both sexual functions in shades of purple. Significant treatment effects, as identified by likelihood ratio tests (Table S6), are indicated by p-values from t-tests conducted for pairwise comparisons between treatment levels shown by black connecting lines.

**Fig. S5.**
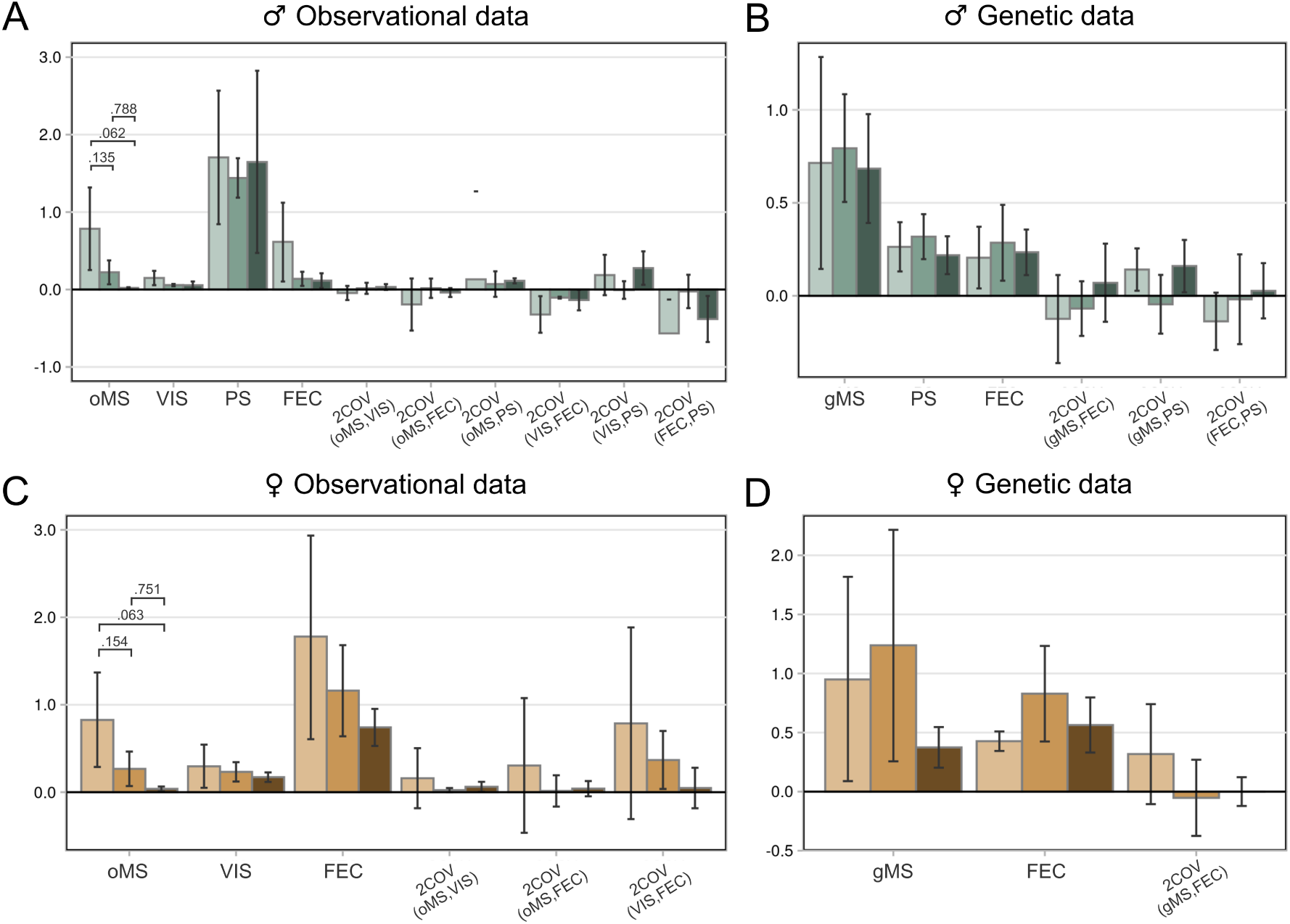
Effect of pollinator abundance treatment (low: light color, medium: medium color, high: dark color) on variance components of outcrossed reproductive success (mean ± 95% bootstrapped confidence intervals), including covariances, in males (A, B) and females (C, D). The analysis focuses on the genetic number of mates (A, C) and the observational number of mates (B, D). When variance components differed significantly between pollinator treatments, as indicated by likelihood ratio tests on nested models (Table S3), adjusted p-values from pairwise comparisons using post-hoc Tukey’s HSD tests are displayed by black connecting lines. Abbreviations: (A) gMS: genetic number of mates, PS: proportion of fertilized seeds per mate, FEC: number of seeds produced per mate, (B) oMS: observational number of mates, VIS: number of visited flowers per mate, PS: proportion of fertilized seeds per mate, FEC: number of seeds produced per mate per visited flowers (C) gMS: genetic number of mates, FEC: number of seeds fertilized per mate, (D) oMS: observational number of mates, FEC: number of seeds fertilized per mate per visited flower, VIS: number of visited flowers per mate.

**Table S1.**
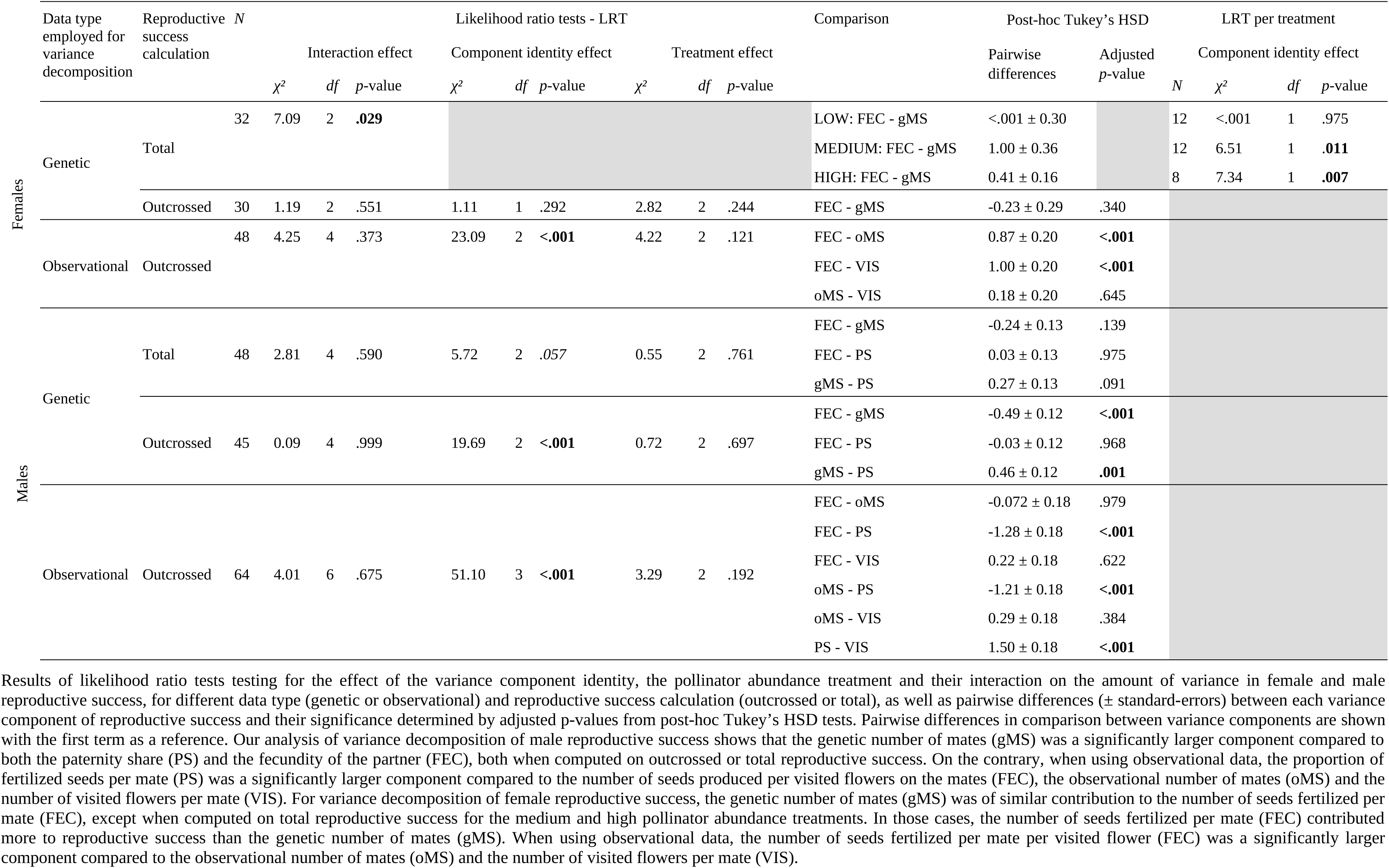
Results of likelihood ratio tests testing for the main component of reproductive successes for both sexual functions according to the data type employed for variance decomposition.

**Table S2.** Effects of pollinator abundance treatment, sex (when appropriate) and their interaction on estimated Bateman gradients.

|  | Variable | N | Likelihood ratio tests |  |  |  |  |  |  |  |  |
| --- | --- | --- | --- | --- | --- | --- | --- | --- | --- | --- | --- |
|  |  |  | Interaction effect |  |  | Sex effect |  |  | Treatment effect |  |  |
| | | | $\chi^2$ | df | p-value | $\chi^2$ | df | p-value | $\chi^2$ | df | p-value |
| Bateman gradients based on total reproductive success | Observational number of mates | 250 | 0.46 | 2 | .793 | 8.44 | 1 | **.004**<sup>x</sup> $\in [7, \infty[$ | 5.35 | 2 | .069 |
|  | Genetic number of mates | 268 | 2.64 | 2 | .267 | 6.58 | 1 | **.010** | 3.63 | 2 | .163 |
|  | Number of flowers visited on mates | 250 | 2.47 | 2 | .290 | 0.09 | 1 | .925 | 5.26 | 2 | .072 |
|  | Number of flowers visited on the plant | 272 | 0.16 | 2 | .922 | 0.02 | 1 | .883 | 5.99 | 2 | **.050** |
|  | Total visit number | 272 | 0.35 | 2 | .841 | <.01 | 1 | .970 | 6.69 | 2 | **.035** |
|  | Average pollen load * | 137 |  |  |  |  |  |  | 0.94 | 2 | .624 |
| Bateman gradients based on outcrossed reproductive success | Observational number of mates | 220 | 0.45 | 2 | .800 | 5.17 | 1 | .134 | 7.84 | 2 | **.020**<sup>x</sup> $\in [8, 17]$ |
|  | Genetic number of mates | 228 | 1.19 | 2 | .552 | 2.08 | 1 | .149 | 6.67 | 2 | **.036** |
|  | Number of flowers visited on mates | 220 | 5.17 | 2 | .075 | 0.02 | 1 | .880 | 5.82 | 2 | .055 |
|  | Number of flowers visited on the plant | 228 | 0.45 | 2 | .800 | 0.84 | 1 | .359 | 6.91 | 2 | **.032** |
|  | Total visit number | 220 | 0.47 | 2 | .800 | 0.11 | 1 | .741 | 5.99 | 2 | **.050** |
|  | Average pollen load * | 113 |  |  |  |  |  |  | 1.74 | 2 | .420 |

Results of likelihood ratio tests are shown with sample size ( $N$ ), chi-squared statistic ( $\chi^2$ ), degrees of freedom ( $df$ ) and associated $p$ -value. For variables relying on the computation of pollen carry-overs (defined as the successive number of visited flowers for which pollen export/import remains effective), the range of carry-over values $x$ for which the result remains significant is specified in the form $x \in [a, b]$ , where $a$ and $b$ denote, respectively, the shortest and longest pollen dispersal kernels for which results remained significant.
\* Results for Bateman gradients based on pollen loads are presented only for female reproductive success, as the amount of pollen deposited on stigmas can influence seed production solely via the female function.

**Table S3.**
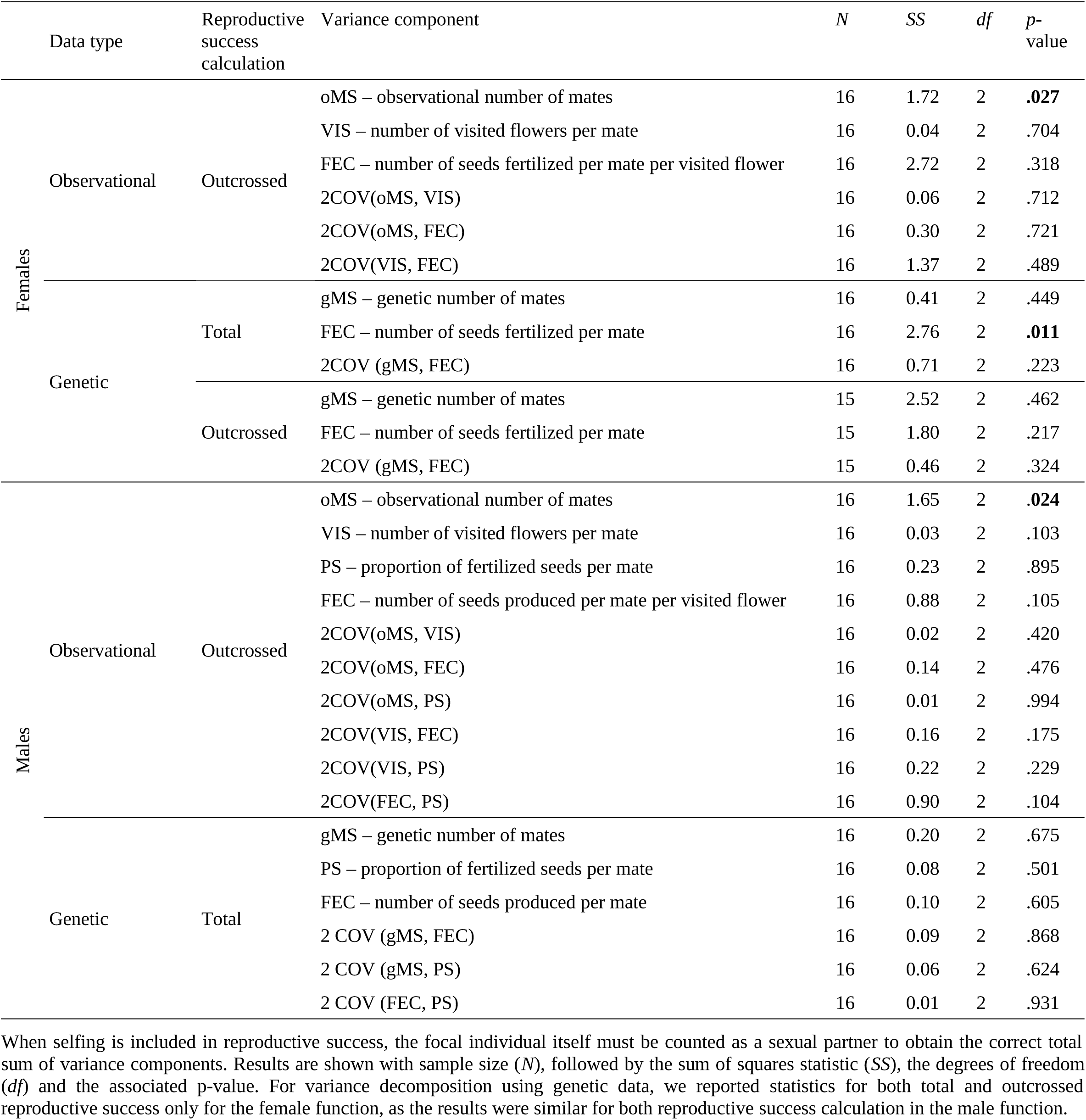
Results of likelihood ratio tests testing for the effect of pollinator abundance treatment on the variance components of the female and male reproductive successes.

**Table S4.** Relationships between male and female fitness components assessed for the three tested pollinator abundances separately.

|  | LOW |  |  |  |  | MEDIUM |  |  |  |  | HIGH |  |  |  |  |
| --- | --- | --- | --- | --- | --- | --- | --- | --- | --- | --- | --- | --- | --- | --- | --- |
| | Estimate | *N* | $\chi^2$ | *df* | *p*-value | Estimate | *N* | $\chi^2$ | *df* | *p*-value | Estimate | *N* | $\chi^2$ | *df* | *p*-value |
| Observational number of mates | 0.04 ± 0.16 | 44 | 0.07 | 1 | .800 | 0.51 ± 0.14 | 54 | 12.21 | 1 | <.001<sup>x</sup> ∈ [1.12] | 0.52 ± 0.18 | 40 | 7.71 | 1 | 0.005<sup>x</sup> ∈ [1.12] |
| Genetic number of mates | 0.10 ± 0.13 | 42 | 0.62 | 1 | .432 | 0.17 ± 0.16 | 50 | 1.17 | 1 | .280 | 0.05 ± 0.11 | 33 | 0.20 | 1 | .658 |
| Outcrossed reproductive success | -0.07 ± 0.26 | 31 | 0.08 | 1 | .773 | -0.20 ± 0.16 | 37 | 1.59 | 1 | .207 | -0.23 ± 0.18 | 29 | 1.74 | 1 | .186 |

Regressions (estimates ± standard-errors) for outcrossed reproductive success, observational and genetic number of mates between male and female sexual function computed for each pollinator abundance. Linear mixed-effects models were used for the regression of male-function values on female-function values, with a random effect of the experimental population. Results of likelihood ratio tests on the female function values are shown with sample size (*N*), chi-squared statistic ( $\chi^2$ ), degrees of freedom (*df*) and associated *p*-value. For variables relying on the computation of pollen carry-overs (defined as the successive number of visited flowers for which pollen export/import remains effective), the range of carry-over values *x* for which the result remains significant is specified in the form *x* ∈ [a,b], where a and b denote, respectively, the shortest and longest pollen dispersal kernels for which results remained significant.

**Table S5.** Results of likelihood ratio tests testing for the effects of pollinator abundance treatment, sex, and their interaction, on the measured variables.

| Variables | Distribution | N | Likelihood ratio tests |  |  |  |  |  |  |  |  |
| --- | --- | --- | --- | --- | --- | --- | --- | --- | --- | --- | --- |
|  |  |  | Treatment effect |  |  | Sex effect |  |  | Interaction effect |  |  |
| | | | $\chi^2$ | df | p-value | $\chi^2$ | df | p-value | $\chi^2$ | df | p-value |
| Total reproductive success <sup>(1)</sup> | Poisson | 270 | 2.49 | 2 | .288 | 0.17 | 1 | .681 | 0.46 | 2 | .793 |
| Outcrossed reproductive success <sup>(1)</sup> | Poisson | 228 | 3.14 | 2 | .208 | 0.02 | 1 | .902 | 0.19 | 2 | .911 |
| Observational number of mates | Poisson | 276 | 8.66 | 2 | **.013**<sup>x ∈ [1.50]</sup> | 0.00 | 1 | 1.000 <sup>(2)</sup> | 0.00 | 2 | 1.000 <sup>(2)</sup> |
| Genetic number of mates | Poisson | 268 | 3.35 | 2 | .187 | 0.05 | 1 | .816 | 0.04 | 2 | .981 |
| Number of flowers visited on mates | Gaussian | 272 | 4.17 | 2 | .124 | 1.75 | 1 | .186 | 0.11 | 2 | .948 |
| Number of flowers visited on the plant | Poisson | 160 | 5.78 | 2 | .056 |  |  |  |  |  |  |
| Total number of visits on the plant | Poisson | 160 | 8.97 | 2 | **.011** |  |  |  |  |  |  |
| Average pollen load on the plant | Poisson | 379 | 1.59 | 2 | .451 |  |  |  |  |  |  |

Results of likelihood ratio tests are shown with sample size (*N*), chi-squared statistic ( $\chi^2$ ), degrees of freedom (*df*) and associated *p*-value. For variables that involved the computation of pollen carry-over (defined as the number of successive flowers visited during which pollen export/import remains effective), the set of carry-over values *x* for which the result remains significant is specified in the form *x* ∈ [*a*,*b*], where *a* and *b* denote, respectively, the shortest and longest pollen dispersal kernels for which results remained significant.
<sup>(1)</sup> Including non-reproducing individuals did not alter the results. There was no effect of sex, treatment, or their interaction on either total or outcrossed reproductive success.
<sup>(2)</sup> The observational number of mates is, on average, identical between the male and female functions, resulting in an absolute null effect of sex and its interaction with treatment. This was expected because the directionality of pollinator visits moving from one plant to the next necessarily leads to symmetry in observed numbers of sexual partners obtained by the male or female function, with the notable exception of the first visit in the sequence (i.e. effective for the male function only) and the last visit (i.e. effective for the female function only). In our analysis, the effects of the first and last visits are minimal because the first and last plants visited by a pollinator during observation sessions were always revisited at least once during the sequence. This finding holds true for all carry-over values explored, ranging from 1 to infinity.

**Table S6.**
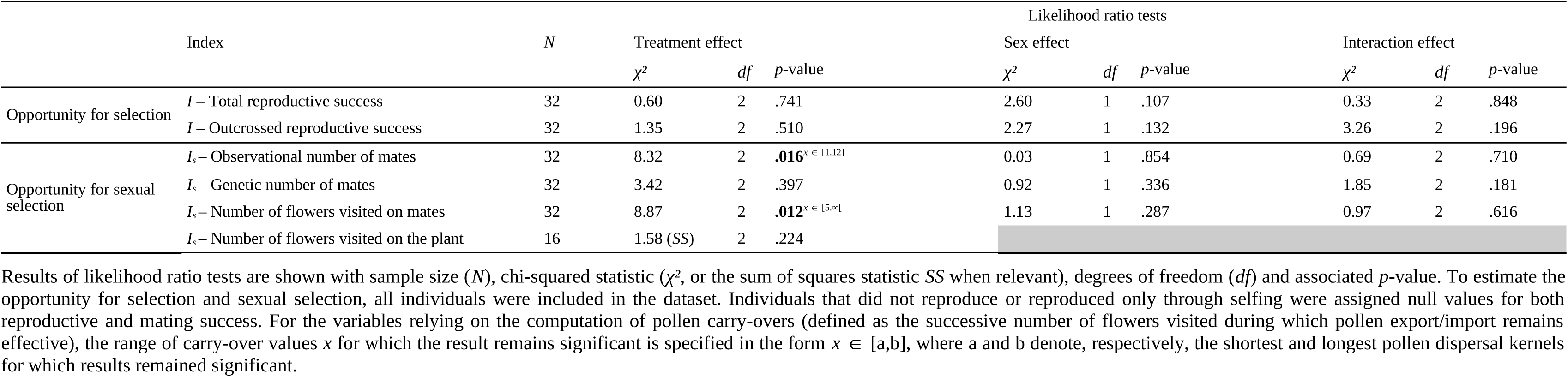
Results of likelihood ratio tests on the effects of sex, treatment, and their interaction on the opportunities for selection ( *I*) and the opportunities for sexual selection (*I_s_*)

**Table S7.** Results of likelihood ratio tests for model comparison on the relationship between genetic and observational number of mates using reproductive skew measures (*Q_gen_* vs. *Q_obs_*)

| Response | Fixed effect | Heteroscedasticity | Test | $\chi^2$ | p-value | Conclusion |
| --- | --- | --- | --- | --- | --- | --- |
| (1) $Q_{gen}$ | $Q_{obs} * \text{Treatment}$ | Yes | | | | |
| (2) $Q_{gen}$ | $Q_{obs} * \text{Treatment}$ | No | (2) vs (1) | 16.65 | <.001 | High heteroscedasticity |
| (3) $Q_{gen}$ | $Q_{obs} + \text{Treatment}$ | Yes | (3) vs (1) | 0.24 | .885 | Similar slopes between treatment modalities |
| (4) $Q_{gen}$ | $Q_{obs}$ | Yes | (4) vs (3) | 1.39 | .845 | Similar intercepts between treatment modalities |
| (5) $Q_{gen}$ | Intercept only | Yes | (5) vs (4) | 11.39 | <.001 | $Q_{obs}$ slope differ from 1 |
| (6) $Q_{gen}$ | Regression line forced through point $Q_{obs} = Q_{gen}$ for largest observed $Q_{obs}$. | Yes | (6) vs (4) | 10.56 | <.01 | $Q_{gen} > Q_{obs}$ for high $Q_{obs}$ and conversely for low $Q_{obs}$. |

The response variable and the fixed effects included in the linear mixed-effect models are indicated. Heteroscedasticity of residuals was accounted for (Yes) or not (No) by fitting the residual variance as a function of the number of genotyped seeds from which each $Q_{gen}$ value was obtained. Likelihood ratio test results comparing models are reported, including the chi-squared statistic ( $\chi^2$ ), associated p-value, and a verbal interpretation of each result.

**Table S8.**
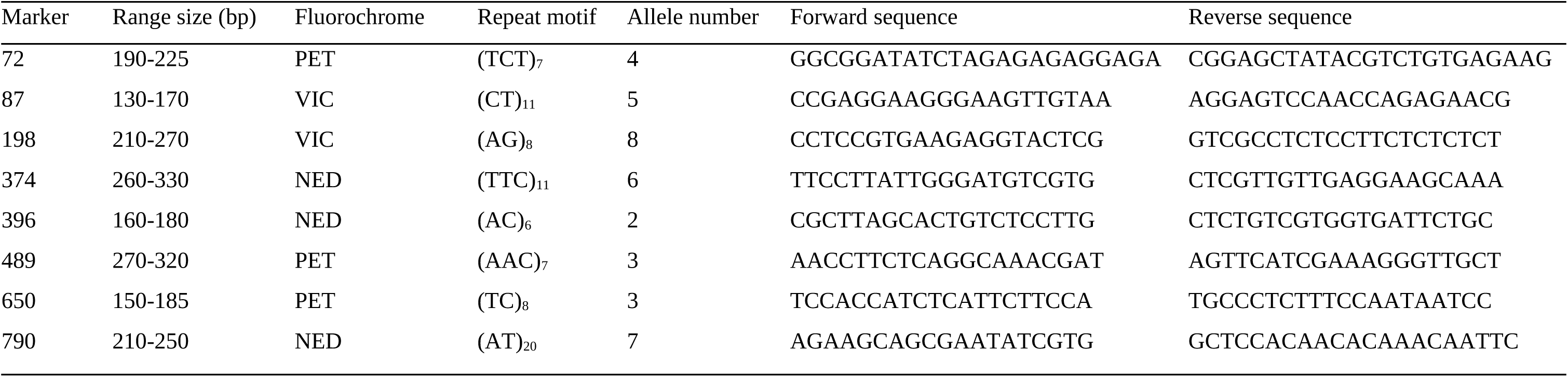
Properties of the eight nuclear microsatellites used for individual allelic characterization and paternity analysis.

**Table S9.**
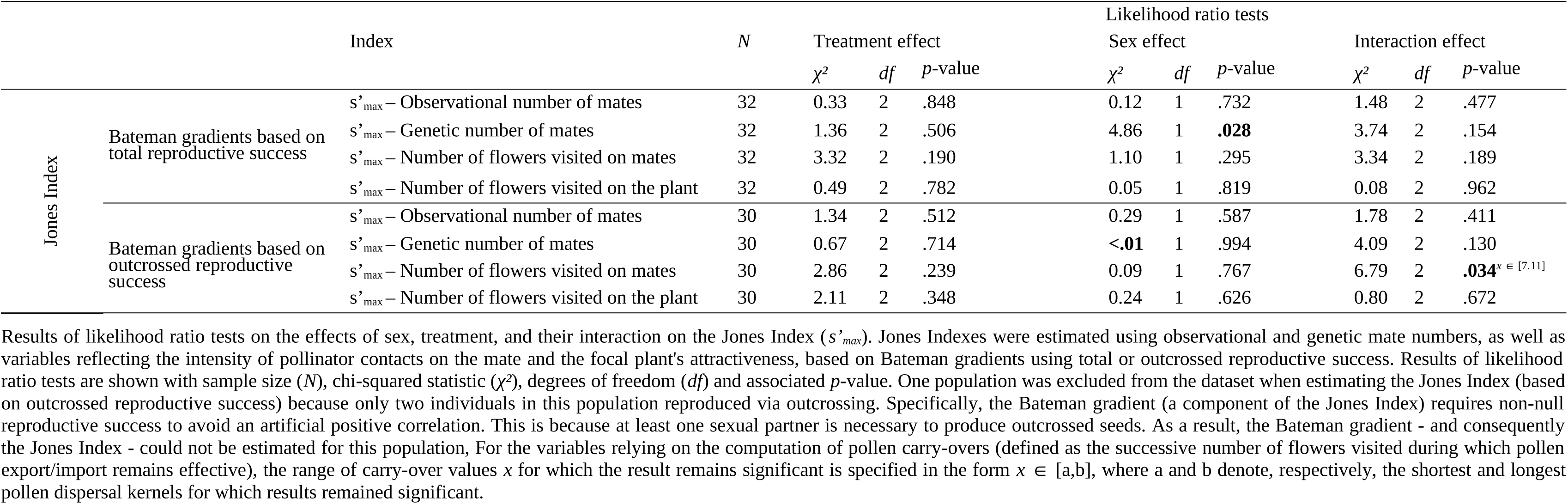
Results of likelihood ratio tests testing for the effects of pollinator abundance treatment, sex, and their interaction, on Jones indexes.

**Table S10.** Results of likelihood ratio tests testing for the cross-sex effects for the Bateman gradients using either the genetic number of mates or the observational number of mates, for each sexual function.

|  |  | LOW |  |  |  | MEDIUM |  |  |  | HIGH |  |  |  |
| --- | --- | --- | --- | --- | --- | --- | --- | --- | --- | --- | --- | --- | --- |
| | | *N* | $\chi^2$ | *df* | *p*-value | *N* | $\chi^2$ | *df* | *p*-value | *N* | $\chi^2$ | *df* | *p*-value |
| Observational number of mates | Female | 35 | 0.32 | 1 | .574 | 42 | 0.37 | 1 | .542 | 34 | 0.01 | 1 | .913 |
|  | Male | 34 | 0.46 | 1 | .498 | 43 | 0.95 | 1 | .330 | 32 | 0.45 | 1 | .500 |
| Genetic number of mates | Female | 33 | 0.52 | 1 | .472 | 40 | 0.14 | 1 | .708 | 32 | 0.06 | 1 | .803 |
|  | Male | 35 | 2.74 | 1 | .098 | 41 | 1.70 | 1 | .192 | 29 | 1.26 | 1 | .261 |

Cross-sex effects were explored for each sex and treatment separately, the regression of outcrossed reproductive success of the focal sex on both (i) the number of mates (either observational or genetic) of the focal sex and (ii) the number of mates (either observational or genetic) of the opposite sex. The latter term represents what is referred to as the cross-sex effect (71). Results of likelihood ratio tests on the cross-sex effect are shown with sample size (*N*), chi-squared statistic ( $\chi^2$ ), degrees of freedom (*df*) and associated *p*-value.

## Notes

### Competing Interest Statement

The authors have declared no competing interest.

## References

1. M. Andersson, Sexual Selection (Princeton University Press, 1994).

2. M. D. Jennions, H. Kokko, “Sexual selection” in Evolutionary Behavioral Ecology (Oxford University Press, Oxford, 2010).

3. T. Janicke, I. K. Häderer, M. J. Lajeunesse, N. Anthes, Darwinian sex roles confirmed across the animal kingdom. Science Advances 2, e1500983 (2016).

4. L. Winkler, M. Moiron, E. H. Morrow, T. Janicke, Stronger net selection on males across animals. eLife 10, e68316 (2021).

5. C. Darwin, The Descent of Man, and Selection in Relation to Sex (John Murray, London, 1871).

6. A. Parker, Sperm competition and its evolutionary consequences in the insects. Biological Reviews 7, 525–567 (1970).

7. G. A. Parker, Conceptual developments in sperm competition: a very brief synopsis. Philosophical Transactions of the Royal Society B: Biological Sciences 375, 20200061 (2020).

8. W. G. Eberhard, Postcopulatory sexual selection: Darwin’s omission and its consequences. Proceedings of the National Academy of Sciences 106, 10025–10032 (2009).

9. T. R. Birkhead, How stupid not to have thought of that: post-copulatory sexual selection. Journal of Zoology 281, 78–93 (2010).

10. J. C. Moore, J. R. Pannell, Sexual selection in plants. Current Biology 21, R176–R182 (2011).

11. P. Pollo, M. Lagisz, Y. Yang, A. Culina, S. Nakagawa, Synthesis of sexual selection: a systematic map of meta-analyses with bibliometric analysis. Biological Reviews 99, 2134– 2175 (2024).

12. A. J. Bateman, Intra-sexual selection in *Drosophila*. Heredity 2, 349–368 (1948).

13. S. J. Arnold, Is there a unifying concept of sexual selection that applies to both plants and animals? The American Naturalist 144, S1–S12 (1994).

14. J. Lehtonen, Bateman gradients from first principles. Nat Commun 13, 3591 (2022).

15. S. Fromonteil, L. Marie-Orleach, L. Winkler, T. Janicke, Sexual selection in females and the evolution of polyandry. PLOS Biology 21, e3001916 (2023).

16. J. Tonnabel, P. David, J. R. Pannell, Do metrics of sexual selection conform to Bateman’s principles in a wind-pollinated plant? Proceedings of the Royal Society B: Biological Sciences 286, 20190532 (2019).

17. E. Barbot, M. Dufaÿ, I. De Cauwer, Sex-specific selection patterns in a dioecious insect-pollinated plant. Evolution 77, 1578–1590 (2023).

18. J. Ollerton, R. Winfree, S. Tarrant, How many flowering plants are pollinated by animals? Oikos 120, 321–326 (2011).

19. C. G. Eckert, S. Kalisz, M. A. Geber, R. Sargent, E. Elle, P.-O. Cheptou, C. Goodwillie, M. O. Johnston, J. K. Kelly, D. A. Moeller, E. Porcher, R. H. Ree, M. Vallejo-Marín, A. A. Winn, Plant mating systems in a changing world. Trends in Ecology & Evolution 25, 35–43 (2010).

20. D. D. L. Gervasi, F. P. Schiestl, Real-time divergent evolution in plants driven by pollinators. Nature Communications 8, 14691 (2017).

21. C. M. Caruso, K. E. Eisen, R. A. Martin, N. Sletvold, A meta-analysis of the agents of selection on floral traits. Evolution 73, 4–14 (2019).

22. S. E. Ramos, F. P. Schiestl, Rapid plant evolution driven by the interaction of pollination and herbivory. Science 364, 193–196 (2019).

23. P.-O. Cheptou, E. Imbert, M. Thomann, Rapid evolution of selfing syndrome traits in *Viola arvensis* revealed by resurrection ecology. American Journal of Botany 109, 1838–1846 (2022).

24. H. Kokko, D. J. Rankin, Lonely hearts or sex in the city? Density-dependent effects in mating systems. Philosophical Transactions of the Royal Society B: Biological Sciences 361, 319–334 (2006).

25. T. Aronsen, A. Berglund, K. B. Mobley, I. I. Ratikainen, G. Rosenqvist, Sex ratio and density affect sexual selection in a sex-role reversed fish: sexual selection in a sex-role reversed fish. Evolution 67, 3243–3257 (2013).

26. E. L. McCullough, B. A. Buzatto, L. W. Simmons, Population density mediates the interaction between pre- and postmating sexual selection. Evolution 72, 893–905 (2018).

27. R. D. Cothran, D. Schmidenberg, A. R. Stiff, G. A. Wellborn, R. A. Relyea, Effects of density on the strength of sexual selection in the laboratory and in nature. Biological Journal of the Linnean Society 140, 504–517 (2023).

28. L. Winkler, R. Eilhardt, T. Janicke, Population density affects sexual selection in an insect model. Functional Ecology 37, 2734–2747 (2023).

29. A. De Nardo, B. Biswas, J. Perdigón Ferreira, A. Meena, S. Lüpold, Socio-ecological context modulates the significance of territorial contest competition in Drosophila prolongata. Proceedings B 292 (2025).

30. N. Sletvold, The context dependence of pollinator-mediated selection in natural populations. International Journal of Plant Sciences 180, 934–943 (2019).

31. D. A. Christopher, R. J. Mitchell, J. D. Karron, Pollination intensity and paternity in flowering plants. Annals of Botany 125, 1–9 (2020).

32. A. Kwok, M. Dorken, Sexual selection on male but not female function in monoecious and dioecious populations of broadleaf arrowhead (Sagittaria latifolia). Proceedings of the Royal Society B: Biological Sciences 289 (2022).

33. M. Hou, Ø. H. Opedal, Z.-G. Zhao, Sexually concordant selection on floral traits despite greater opportunity for selection through male fitness. New Phytologist 241, 926–936 (2024).

34. Z. Gauthey, C. Tentelier, O. Lepais, A. Elosegi, L. Royer, S. Glise, J. Labonne, With our powers combined: integrating behavioral and genetic data to estimate mating success and sexual selection. Rethinking Ecology 2, 1–26 (2017).

35. F. R. Badenes-Pérez, Benefits of insect pollination in Brassicaceae: a meta-analysis of self-compatible and self-incompatible crop species. Agriculture 12, 446 (2022).

36. R. Rader, B. G. Howlett, S. A. Cunningham, D. A. Westcott, L. E. Newstrom-Lloyd, M. K. Walker, D. A. J. Teulon, W. Edwards, Alternative pollinator taxa are equally efficient but not as effective as the honeybee in a mass flowering crop. Journal of Applied Ecology 46, 1080–1087 (2009).

37. M. J. Wade, Sexual selection and variance in reproductive success. The American Naturalist 114, 742–764 (1979).

38. M. J. Wade, S. J. Arnold, The intensity of sexual selection in relation to male sexual behaviour, female choice, and sperm precedence. Animal Behaviour 28, 446–461 (1980).

39. S. J. Arnold, Bateman’s principles and the measurement of sexual selection in plants and animals. The American Naturalist 144, S126–S149 (1994).

40. A. G. Jones, On the opportunity for sexual selection, the Bateman gradient and the maximum intensity of sexual selection. Evolution 63, 1673–1684 (2009).

41. J. M. Henshaw, A. T. Kahn, K. Fritzsche, A rigorous comparison of sexual selection indexes via simulations of diverse mating systems. Proceedings of the National Academy of Sciences 113, E300–E308 (2016).

42. B. Pélissié, P. Jarne, V. Sarda, P. David, Disentangling precopulatory and postcopulatory sexual selection in polyandrous species. Evolution 68, 1320–1331 (2014).

43. S. Lüpold, L. Jin, W. B. Liao, Population density and structure drive differential investment in pre- and postmating sexual traits in frogs. Evolution 71, 1686–1699 (2017).

44. N. Anthes, I. K. Häderer, N. K. Michiels, T. Janicke, Measuring and interpreting sexual selection metrics: evaluation and guidelines. Methods in Ecology and Evolution 8, 918–931 (2016).

45. L. Rowe, D. Houle, The lek paradox and the capture of genetic variance by condition dependent traits. Proceedings of the Royal Society of London. Series B: Biological Sciences 263, 1415–1421 (1996).

46. L. R. Dougherty, Meta-analysis reveals that animal sexual signalling behaviour is honest and resource based. Nat Ecol Evol 5, 688–699 (2021).

47. A. Roulin, Condition-dependence, pleiotropy and the handicap principle of sexual selection in melanin-based colouration. Biol Rev Camb Philos Soc 91, 328–348 (2016).

48. S. Cotton, J. Small, A. Pomiankowski, Sexual selection and condition-dependent mate preferences. Current Biology 16, R755–R765 (2006).

49. A. J. Lumley, Ł. Michalczyk, J. J. N. Kitson, L. G. Spurgin, C. A. Morrison, J. L. Godwin, M. E. Dickinson, O. Y. Martin, B. C. Emerson, T. Chapman, M. J. G. Gage, Sexual selection protects against extinction. Nature 522, 470–473 (2015).

50. J. G. Cally, D. Stuart-Fox, L. Holman, Meta-analytic evidence that sexual selection improves population fitness. Nat Commun 10, 2017 (2019).

51. E. Noël, E. Fruitet, D. Lelaurin, N. Bonel, A. Ségard, V. Sarda, P. Jarne, P. David, Sexual selection and inbreeding: Two efficient ways to limit the accumulation of deleterious mutations. Evol Lett 3, 80–92 (2018).

52. M. Almbro, L. W. Simmons, Sexual selection can remove an experimentally induced mutation load. Evolution 68, 295–300 (2014).

53. J. M. Parrett, S. Chmielewski, E. Aydogdu, A. Łukasiewicz, S. Rombauts, A. Szubert-Kruszyńska, W. Babik, M. Konczal, J. Radwan, Genomic evidence that a sexually selected trait captures genome-wide variation and facilitates the purging of genetic load. Nat Ecol Evol 6, 1330–1342 (2022).

54. J. Tonnabel, P. David, T. Janicke, A. Lehner, J.-C. Mollet, J. R. Pannell, M. Dufay, The scope for postmating sexual selection in plants. Trends Ecol Evol 36, 556–567 (2021).

55. S. Takayama, A. Isogai, Self-incompatibility in plants. Annu Rev Plant Biol 56, 467–489 (2005).

56. J. B. Nasrallah, Plant mating systems: self-incompatibility and evolutionary transitions to self-fertility in the mustard family. Curr Opin Genet Dev 47, 54–60 (2017).

57. P. H. Williams, C. B. Hill, Rapid-cycling populations of *Brassica*. Science 232, 1385–1389 (1986).

58. A. Dafni, P. Kevan, C. L. Gross, K. Goka, *Bombus terrestris*, pollinator, invasive and pest: an assessment of problems associated with its widespread introductions for commercial purposes. Appl. Entomol. Zool. 45, 101–113 (2010).

59. L. Chittka, J. D. Thomson, “Behavioral responses of pollinators to variation in floral display size and their influences on the evolution of floral traits” in Cognitive Ecology of Pollination - Animal Behavior and Floral Evolution (Cambridge University Press, Cambridge, 2001).

60. O. Friard, M. Gamba, BORIS: a free, versatile open-source event-logging software for video/audio coding and live observations. Methods in Ecology and Evolution 7, 1325–1330 (2016).

61. N. Anthes, P. David, J. R. Auld, J. N. A. Hoffer, P. Jarne, J. M. Koene, H. Kokko, M. C. Lorenzi, B. Pélissié, D. Sprenger, A. Staikou, L. Schärer, Bateman gradients in hermaphrodites: an extended approach to quantify sexual selection. The American Naturalist 176, 249–263 (2010).

62. L. S. Adler, R. E. Irwin, Comparison of pollen transfer dynamics by multiple floral visitors: experiments with pollen and fluorescent dye. Ann Bot 97, 141–150 (2006).

63. C. Minnaar, B. Anderson, Using quantum dots as pollen labels to track the fates of individual pollen grains. Methods in Ecology and Evolution 10, 604–614 (2019).

64. N. M. Waser, M. V. Price, Experimental studies of pollen carryover: effects of floral variability in *Ipomopsis aggregata*. Oecologia 62, 262–268 (1984).

65. M. C. Castellanos, P. Wilson, J. D. Thomson, Pollen transfer by hummingbirds and bumblebees, and the divergence of pollination modes in penstemon. Evolution 57, 2742– 2752 (2003).

66. J. E. Cresswell, A method for quantifying the gene flow that results from a single bumblebee visit using transgenic oilseed rape,Brassica napus L. cv. Westar. Transgenic Research 3, 134–137 (1994).

67. C. Minnaar, B. Anderson, M. L. de Jager, J. D. Karron, Plant–pollinator interactions along the pathway to paternity. Annals of Botany 123, 225–245 (2019).

68. G. Ne’eman, A. Jürgens, L. Newstrom-Lloyd, S. G. Potts, A. Dafni, A framework for comparing pollinator performance: effectiveness and efficiency. Biological Reviews 85, 435–451 (2010).

69. T. C. Marshall, J. Slate, L. E. B. Kruuk, J. M. Pemberton, Statistical confidence for likelihood-based paternity inference in natural populations. Molecular Ecology 7, 639–655 (1998).

70. S. T. Kalinowski, M. L. Taper, T. C. Marshall, Revising how the computer program Cervus accommodates genotyping error increases success in paternity assignment. Molecular Ecology 16, 1099–1106 (2007).

71. B. Pélissié, P. Jarne, P. David, Sexual selection without sexual dimorphism: Bateman gradients in a simultaneous hermaphrodite. Evolution 66, 66–81 (2012).

72. L. Marie-Orleach, T. Janicke, D. B. Vizoso, P. David, L. Schärer, Quantifying episodes of sexual selection: Insights from a transparent worm with fluorescent sperm. Evolution 70, 314–328 (2016).

73. S. P. De Lisle, E. I. Svensson, On the standardization of fitness and traits in comparative studies of phenotypic selection. Evolution 71, 2313–2326 (2017).

74. K. Tsuji, E. Kasuya, What do the indices of reproductive skew measure? The American naturalist 158, 155–65 (2001).

75. M. Morisita, *I*_\textrmD-index, a measure of dispersion of individuals. Res. Popul. Ecol., 1– 7 (1962).

76. F. Rousset, J.-B. Ferdy, Testing environmental and genetic effects in the presence of spatial autocorrelation. Ecography 37, 781–790 (2014).

## References

1. Alexander MP. Differential staining of aborted and nonaborted pollen. Stain Technol. 1969;44: 117–122. doi:10.3109/10520296909063335

2. Schneider CA, Rasband WS, Eliceiri KW. NIH Image to ImageJ: 25 years of image analysis. Nat Methods. 2012;9: 671–675. doi:10.1038/nmeth.2089

3. Tonnabel J, David P, Pannell JR. Do metrics of sexual selection conform to Bateman’s principles in a wind-pollinated plant? Proc R Soc B Biol Sci. 2019;286: 20190532. doi:10.1098/rspb.2019.0532

4. Winkler L, Eilhardt R, Janicke T. Population density affects sexual selection in an insect model. Funct Ecol. 2023;37: 2734–2747. doi:10.1111/1365-2435.14410

5. Pélissié B, Jarne P, Sarda V, David P. Disentangling precopulatory and postcopulatory sexual selection in polyandrous species. Evolution. 2014;68: 1320–1331. doi:10.1111/evo.12353

